# Ionic liquid-coated lipid nanoparticles demonstrate prolonged circulation and brain uptake via red blood cell hitchhiking

**DOI:** 10.1101/2025.04.16.649225

**Authors:** Purva Khare, Sara X. Edgecomb, Christine M. Hamadani, Duoyi Hu, Balaji Govindaswamy, George R. Taylor, Rocco Caprara, Eden E. L. Tanner, Devika S Manickam

**Affiliations:** Graduate School of Pharmaceutical Sciences, Duquesne University, Pittsburgh, PA 15282; Department of Chemistry and Biochemistry, The University of Mississippi, MS 38677; Department of Neurology, The University of Texas Health Science Center at Houston, TX 77030

**Author notes:** **Corresponding author:**Devika S Manickam 6431 Fannin Street, MSB 7.102 Houston, TX 77030.

**Keywords:** Lipid nanoparticles, LNPs, ionic liquids, choline *trans*-2-hexenoate, red blood cell-hitchhiking, pharmacokinetics, biodistribution, brain delivery

## Abstract

Lipid nanoparticles (**LNPs**) have transformed the delivery of nucleic acid therapeutics; however, their natural tropism favors the liver resulting in clearance by the reticuloendothelial system, with less than 1% of the injected dose reaching challenging targets such as brain. Biocompatible ionic liquids (**ILs**) are tunable materials that can modulate nanoparticle interactions with blood components. Choline *trans*-2-hexenoate (**C2HA**) is an IL known to facilitate red blood cell (**RBC**) hitchhiking of PLGA polymeric nanoparticles and reduces hepatic uptake and therefore enabling transport to distant non-hepatic organs. We wanted to determine if C2HA coatings can show similar RBC hitchhiking effects with LNPs. We previously demonstrated that IL-coated LNPs reduce serum protein binding to LNPs—a key contributor to rapid clearance via the liver. While LNPs coated with choline *trans*-2-hexenoate at 1:1 and 1:2 cation: anion ratios decreased mouse serum protein binding and improved cellular uptake into brain endothelial cells (**BECs**) and motor neurons, they did not show hitchhiking behavior. To identify IL formulations capable of this behavior, we screened higher IL cation: anion ratios (1:3 and 1:5) for LNP coating and optimized IL volumes that allowed stable particle diameters. The resulting IL-coated LNPs successfully hitchhiked on both mouse and human RBCs and significantly enhanced uptake in b.End3 mouse BECs, and NSC-34 neuroblastoma cells compared to uncoated LNPs. 1:3 IL-coated LNPs demonstrated the most pronounced improvement in RBC binding. Whole blood pharmacokinetic and biodistribution analyses demonstrated that IL-coating significantly extends the circulation time of LNPs and results in reduced hepatic uptake alongside increased routing to the brain, compared to standard LNPs. These findings reveal that ILs can be leveraged to re-engineer clinically approved LNP platforms to promote RBC hitchhiking behavior and further be developed for drug delivery to challenging extra-hepatic targets such as the brain.

**Graphical abstract:** 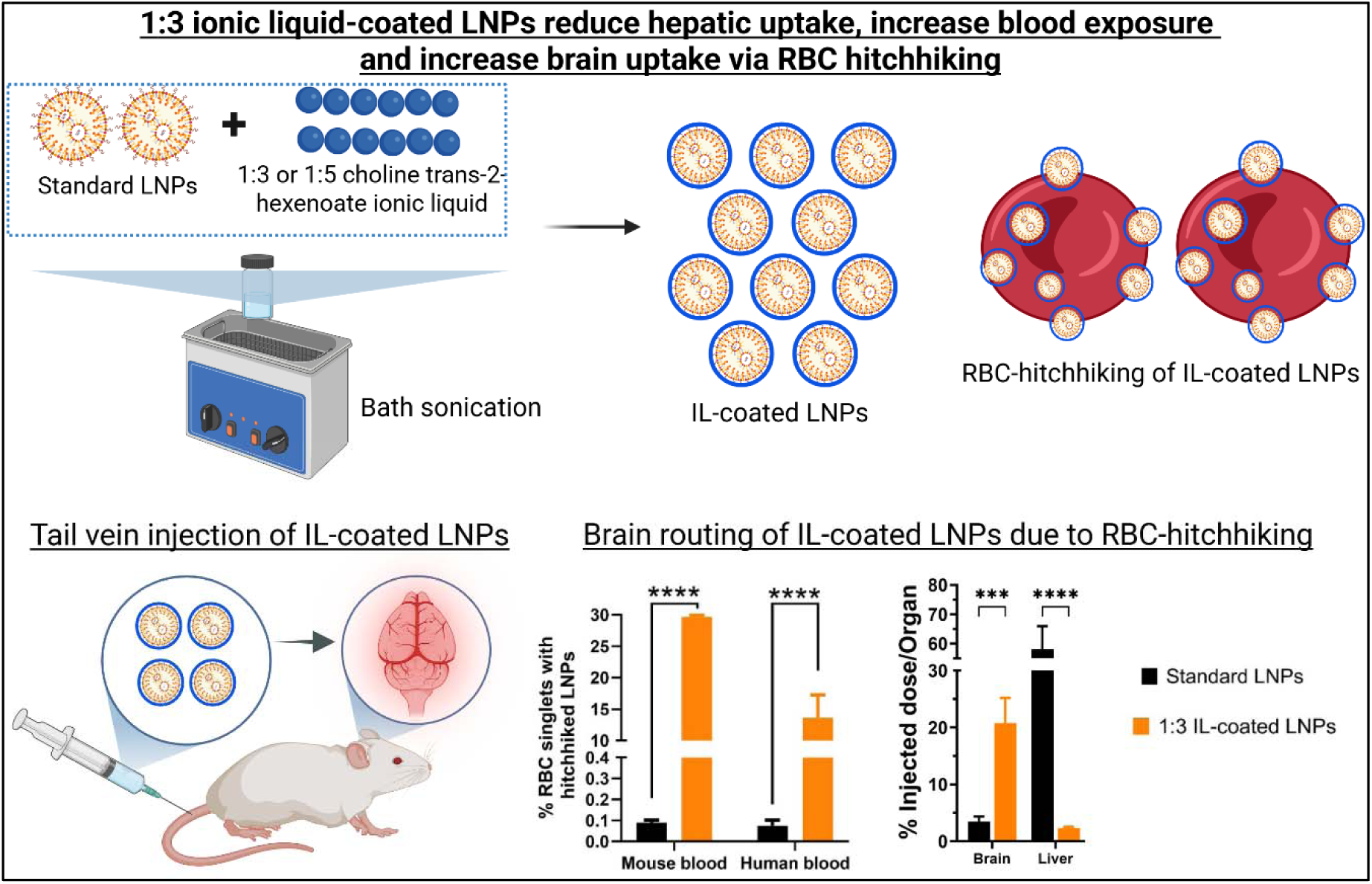

## Introduction

Lipid nanoparticles (**LNPs**) have transformed the field of large molecule drug delivery and are extensively studied for RNA delivery to muscle and liver targets (**Figure 1a**) [1–3]. They are used as nucleic acid carriers in two FDA-approved drugs: Moderna and Pfizer-BioNTech COVID-19 mRNA vaccines and as siRNA carriers in Onpattro [1, 2, 4]. LNPs have reported siRNA loading efficiencies ca. 75-80% [5–7], a significant advantage for drug delivery. The high encapsulation efficiencies of LNPs can be harnessed to improve drug delivery to well-vascularized tissues like the lungs, heart, and the blood-brain barrier (**BBB**)—extra-hepatic tissues that are comparatively challenging to target [8].

**Figure 1.**
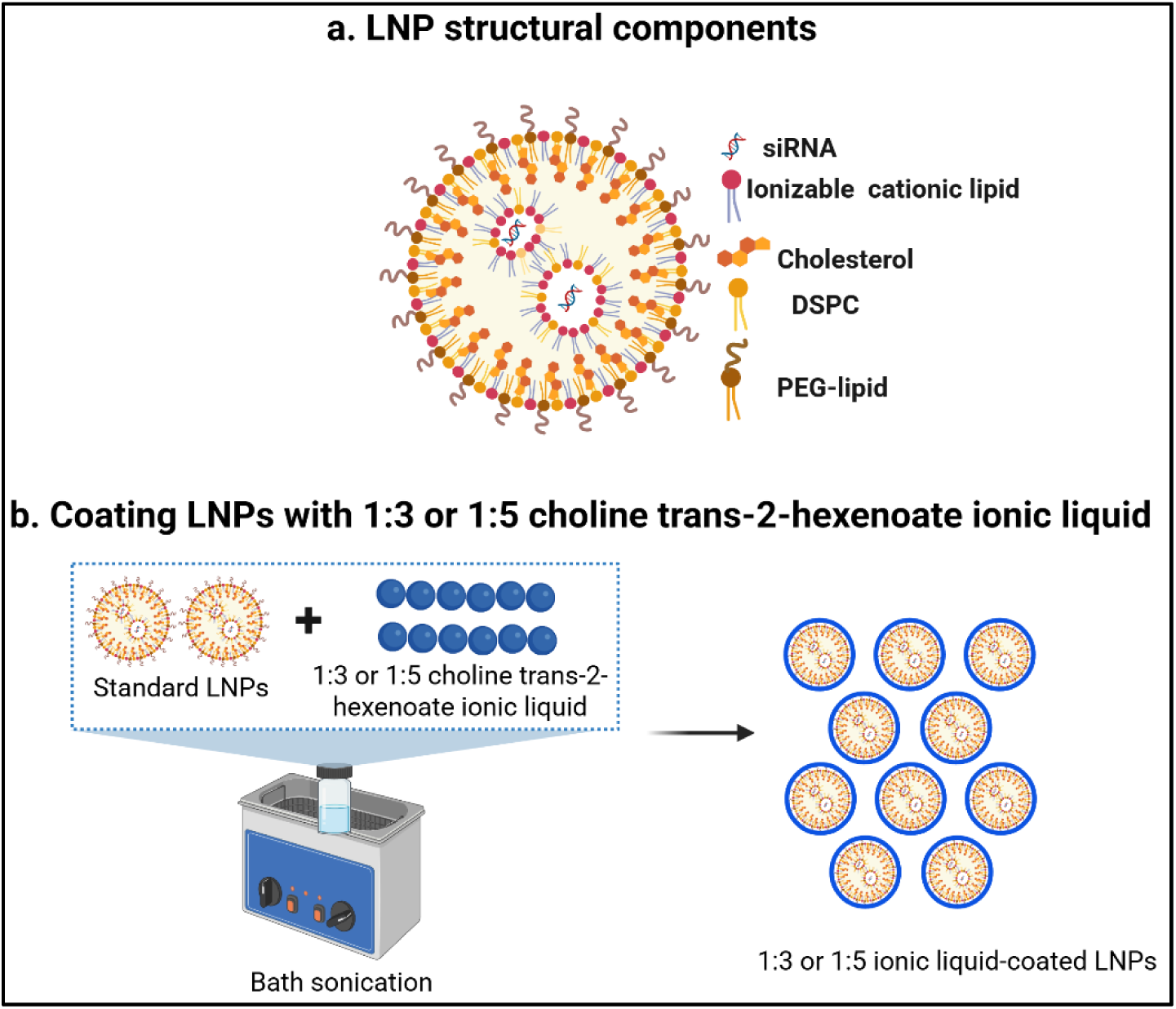
Schematic representation of the structure of siRNA-loaded lipid nanoparticles (LNPs), and the coating process using choline *trans*-2-hexenoate ionic liquid (IL). (**a**) The siRNA-LNP structure highlighting its lipid components: ionizable cationic lipid, cholesterol, DSPC, and PEG-lipid. (**b**) Illustration of the coating process for LNPs with choline *trans*-2-hexenoate IL at 1:3 and 1:5 cation-to-anion ratios (created with Biorender.com).

An example that highlights the importance of targeting distant tissues is the unaddressed clinical need to deliver drugs to brain endothelial cells (**BECs**), which are essential components of the BBB. Dysfunctional/damaged BECs contribute to a range of neurological disorders [9–11]. Targeting BECs for drug delivery presents a promising approach to restoring neuronal function without having to cross the BBB—commonly acknowledged as a holy grail of brain drug delivery. While LNPs have achieved clinical approval, their application for targeting extra-hepatic/distant tissues is still a challenge. A key limitation is their inability to deliver therapeutically meaningful doses to extra-hepatic organs such as the lungs, heart, or brain. One major consideration is related to the established mechanism of LNP uptake by one of the most prominent clearance organs, the liver. Following *i.v.* administration, apolipoprotein E adsorption onto LNP surfaces has been identified as crucial for their uptake into liver cells/hepatocytes [12]. The resulting apolipoprotein E-LNP complex functions as a ligand for lipoprotein receptors on hepatocytes, facilitating their endocytosis and subsequent liver accumulation [13]. Therefore, LNPs are inherently directed to the liver whereas distant organs like the lungs, heart, duodenum, and BBB receive negligible doses [14].

Ionic liquids (**ILs**) are a class of materials consisting of asymmetric and bulky cations and anions [15–17]. When composed of ingredients with established biosafety, the ionic liquids can be used in biological contexts for drug delivery applications. Note that for simplicity we use the term “ionic liquid” throughout, but the materials described herein share properties of both traditional ILs and deep eutectic solvents, since the choline and carboxylic acids are present in non-stoichiometric ratios and likely engage in dynamic protonation. Interactions between the ions of the IL govern the physicochemical properties within the IL and its interaction with the external microenvironment such as blood and plasma proteins. ILs have demonstrated the ability to enhance biocompatibility [18], improve the pharmacokinetic profiles of drug [19–21], and have been used as transfection agents [22, 23]. We propose to utilize and expand the physicochemical properties of ILs to LNPs to regulate the bio-interactions of LNPs in the bloodstream. This can be achieved by coating LNPs with choline *trans*-2-hexenoate IL to modify the surfaces of LNPs (**Figure 1b**).

The choline *trans*-2-hexenoate IL is composed of the choline cation and 2-hexenoic acid anion, both of which are derived from naturally occurring components in the body and are non-immunogenic. Choline *trans*-2-hexenoate also imparts protein-avoidant properties to nanoparticles, which could potentially decrease opsonization and prolong the circulation time of nanoparticles associated with the IL [24]. When coated on poly(lactic-co-glycolic) acid (**PLGA**) nanoparticles, the choline *trans*-2-hexenoate IL facilitated red blood cell (**RBC**) hitchhiking of nanoparticles and decreased clearance 30-fold post-*i.v.* injection in comparison to bare nanoparticles [25, 26]. RBC hitchhiking is a process in which nanoparticles associated with ILs attach to RBC membranes, facilitating their transport to well-vascularized organs such as lungs and brain. RBC hitchhiking is potentially derived from interactions of the 2-hexenoic acid anion with RBC membrane components [24, 27]. Subsequently, accumulation in the target organs occurs as RBCs exit circulation, shearing through blood vessel capillaries and depositing nanoparticles onto endothelial cells [18, 24, 28]. Nanoparticles can consecutively enter the endothelial cells using receptor-mediated endocytosis. Building upon this concept, LNPs coated with ILs may also bind to RBCs resulting in RBC hitchhiking. However, whether this IL can facilitate RBC hitchhiking of soft particles such as LNPs has not previously been explored. Since LNPs have a fundamentally different surface chemistry than PLGA polymeric nanoparticles, it is unclear whether the same IL will enable similar RBC interactions. Moreover, as LNPs are known to interact with blood components in ways that promote clearance, modifying their surface interactions is key to unlocking their potential for targeted delivery to extra-hepatic tissues. We hypothesized that modifying the LNP surface with ILs may regulate their blood interactions in a manner conducive to hitchhiking.

In this pilot study, we developed siRNA-loaded LNPs using C12-200 as the ionizable cationic lipid and then coated pre-formed LNPs with choline *trans*-2-hexenoate IL at two different cation:anion ratios (1:3 and 1:5). The coating and formulation processes were optimized, and the colloidal properties and stability of IL-coated LNPs were characterized using dynamic light scattering. To model cellular uptake in distant, hard-to-reach tissues, we selected two CNS cell lines: mouse brain endothelial cells (**bEnd.3**) and mouse neuroblastoma (**NSC-34**).

While these models do not fully replicate the complexity of the BBB or neuronal networks, they serve as representative systems for evaluating delivery to CNS-associated tissues. Importantly, our intent was not to demonstrate BBB penetration but rather to use these cells as surrogates for assessing whether IL-coated LNPs can be internalized by cell types located in distant tissues. To measure delivery efficiency, we used flow cytometry to quantify the uptake of Cy5-labeled siRNA into both bEnd.3 and NSC-34 cells. Additionally, the hemolysis and RBC hitchhiking potential of the optimized IL-coated LNP formulations were examined using a plate reader and flow cytometry assays. In summary, this pilot study supports the feasibility of using biocompatible ILs like choline *trans*-2-hexenoate to re-engineer LNPs for RBC hitchhiking. By modifying LNP interactions with blood components, especially RBCs, this approach has the potential to shift biodistribution away from clearance organs such as the liver. We tested our hypothesis by conducting whole blood pharmacokinetic and biodistribution studies which showed that IL-coating significantly extended the circulation time of LNPs with reduced hepatic accumulation alongside increased routing to the brain, compared to standard LNPs. Overall, this work provides a foundation for the development of next-generation LNP systems capable of overcoming current barriers for extra-hepatic drug delivery.

## Experimental section

### Materials

1,2-distearoyl-sn-glycero-3-phosphocholine (DSPC) (850365P) and 1,2-dimyristoyl-rac-glycero-3-methoxypolyethylene glycol-2000 (PEG-DMG) (8801518) were obtained from Avanti Polar Lipids (Alabaster, AL). Cholesterol (8667) and Cy5-labeled siRNA were sourced from Sigma-Aldrich (St. Louis, MO), along with choline bicarbonate (80% in water) and *trans*-2-hexenoic acid (99%). The ionizable cationic lipid 1,1‘-((2-(4-(2-((2-(bis(2-hydroxy-dodecyl)amino)ethyl)(2-hydroxydodecyl)amino)ethyl)piperazin-1 yl)ethyl)azanediyl)bis(dodecan-2-ol)) (C12-200) was kindly provided by Dr. Muthiah Manoharan, Alnylam Pharmaceuticals (Cambridge, MA). Phosphate buffered saline (PBS) and heat-inactivated fetal bovine serum (FBS) were purchased from Hyclone Laboratories (Logan, UT), while Penicillin-Streptomycin (Pen-Strep) was procured from Gibco (Grand Island, NY). TrypLE Express and DMEM high glucose were obtained from Life Technologies Corporation (Grand Island, NY). Mouse brain endothelial bEnd.3 cells were purchased from American type cell culture (ATCC) (Manassas, VA), and mouse neuroblastoma NSC-34 cells were acquired from CELLutions Biosystems (Burlington, Canada). siGFP (AM4626) was obtained from Thermo Fisher Scientific (Austin, TX). All reagents were used as supplied unless specified otherwise.

### Ionic liquid synthesis and characterization

To synthesize choline and *trans*-2-hexenoate IL (CA2HA) at a 1:3 or 1:5 molar ionic ratio, choline bicarbonate (80% in water, Sigma Aldrich, # C7519) was combined together as previously described with the respective amount of 98% commercially purified *trans*-2-hexenoic acid anion (Sigma Aldrich, #W316903) in a 500 mL round bottom flask at 40°C and allowed to react overnight [29–31]. To remove volatile impurities, each IL was rotary evaporated at 15 mbar for 2 hours at 60°C and dried in a vacuum oven for 72 hours before physical characterization by Karl Fischer titration and proton Nuclear Magnetic Resonance Spectroscopy (^1^HNMR), as reported below:

CA2HA 1:3 (density 1.28 g/mL, MW 446.59, yield: 92.0%) : 1H NMR (400 MHz, DMSO) δ 6.68 (dt, J = 14.7, 7.3 Hz, 3H), 5.78 (d, J = 15.4 Hz, 3H), 3.89 (d, J = 16.3 Hz, 2H), 3.44 (q, J = 4.5 Hz, 2H), 3.13 (s, 9H), 2.13 (d, J = 7.3 Hz, 6H), 1.46 – 1.40 (m, 6H), 0.90 (t, J = 7.0 Hz, 9H).

CA2HA 1:5 (density 1.14 g/mL, MW 674.87, yield: 87.7%) : 1H NMR (400 MHz, DMSO) δ 6.74 (dd, J = 15.3, 7.6 Hz, 5H), 5.78 (d, J = 15.5 Hz, 5H), 3.92 (s, 2H), 3.43 (q, J = 4.4 Hz, 2H), 3.13 (s, 9H), 2.14 (q, J = 7.7, 7.2 Hz, 10H), 1.43 (p, J = 7.1, 6.4 Hz, 10H), 0.91 (td, J = 8.5, 7.5, 2.7 Hz, 15H).

### Preparation of standard and IL-coated siRNA-LNPs

Uncoated LNPs were prepared following our previously established methods [32–34]. A 1 mg/mL siRNA solution was prepared in 10 mM citrate buffer (pH 4.0). Lipid components— C12-200 (ionizable cationic lipidoid), cholesterol, DSPC, and PEG-DMG—were dissolved in ethanol at a molar ratio of 50:10:38.5:1.5. The formulation scheme is detailed in **Table 1**. The lipid phase was added dropwise to the siRNA solution while vortexing at setting ‘7’ on a Fisher benchtop vortexer for 30 seconds. The final siRNA concentration was adjusted to 400 nM by adding nuclease-free deionized water. The cationic lipidoid/siRNA w/w ratio was maintained at 5:1, and the final siRNA concentration remained 400 nM unless specified otherwise. For coated LNPs, siRNA-LNPs were further treated with 5, 10, or 20 µL of 1:3 or 1:5 (cation: anion) choline *trans*-2-hexenoate IL. LNPs were first diluted with nuclease-free deionized water, transferred to glass scintillation vials, and sonicated in a bath sonicator for 1 minute. Afterwards, the specified volume of IL (5 μL -20 μL) was added, followed by 60 minutes of sonication with an ice pack to prevent overheating of the samples. For *in vivo* studies, Cy5-labeled siRNA was used and LNPs from 4-5 batches were concentrated before tail vein injections into mice.

**Table 1.** Representative formulation scheme for standard siRNA-LNPs

| **Component** | **w/w% in final LNP mixture** | **Stock concentration (mg/mL)** | **Volume of the stock solution required (μL)** |
| --- | --- | --- | --- |
| **Ethanol phase** |  |  |  |
| C12-200 | 50 | 1 | 12.5 |
| Cholesterol | 38.5 | 1 | 3.1 |
| PEG-DMG | 1.5 | 0.1 | 13.6 |
| DSPC | 10 | 0.1 | 16.9 |
| Ethanol | - | - | 78.9 |
| **Total ethanol phase volume** |  |  | 125 |
| **Aqueous phase** |  |  |  |
| siRNA or Cy5 siRNA | 400 nM | 1 | 2.5 |
| 10 mM citrate buffer | - | - | 122.5 |
| Nuclease-free deionized water | - | - | 250 |
| **Total aqueous phase volume** |  |  | 375 |
| **Final volume of LNPs prepared (μL)** |  |  | 500 |

### Dynamic Light Scattering

The physicochemical characteristics of LNPs was studied by measuring average particle diameters, dispersity indices and zeta potentials using dynamic light scattering on a Malvern Zetasizer Pro (Malvern Panalytical Inc., Westborough, PA). When indicated, measurements were conducted over seven days with intermittent storage at 2–8 °C. Standard and IL-coated LNPs were diluted to a final siRNA concentration of 50 nM in nuclease-free water for particle size and zeta potential measurements. All measurements were performed in triplicate, and results are reported as mean ± standard deviation (SD).

### Cell culture

NSC-34 and bEnd.3 cells were grown in DMEM high glucose medium supplemented with 10% FBS and 1% Pen-Strep. The culture medium was refreshed every 48 hours until cells reached 90% confluency. All cell cultures were kept in a humidified incubator at 37 °C flushed with 5% CO_2_.

### Cytocompatibility of LNPs

bEnd.3 and NSC-34 cells were seeded at 16,500 cells/well in Poly-D-Lysine-coated 96-well plates (Azer Scientific, Morgantown, PA). Untreated cells were used as a control. Transfections were performed for 4 hours using standard or IL-coated LNPs containing 50 nM siRNA in 50 µL of complete growth medium per well. After four hours, the transfection mixture was replaced with 200 µL of fresh growth medium, followed by a 24-hour incubation at 37 °C and 5% CO_2_. Relative intracellular ATP levels were assessed 24 hours post-transfection using the Cell Titer-Glo luminescence assay to measure cell viability. Relative ATP levels were compared to untreated controls. To each well, 60 µL of complete growth medium and 60 µL of Cell Titer-Glo 2.0 reagent were added to lyse cells. Plates were incubated for 15 minutes at room temperature in the dark on an incubator shaker (Thermo Fisher Scientific, Waltham, MA). After incubation, 60 µL of the lysate was transferred to a white, opaque 96-well plate (Azer Scientific, Morgantown, PA), and luminescence was measured using a SYNERGY HTX multi-mode reader (BioTek Instruments, Winooski, VT). Relative ATP levels (%) were calculated by normalizing luminescence of treated samples to that of untreated cells using **Eq. 1**.

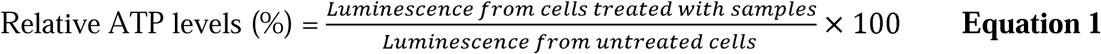

### Cy5 siRNA uptake into b.End3 BECs and NSC-34 neuroblastoma cells using flow cytometry

b.End3 endothelial cells and NSC-34 motor neurons were seeded at 100,000 cells/well in poly-D-Lysine-coated 24-well plates (Genesee Scientific, San Diego, CA) in complete growth medium. The cells were allowed to acclimate in a humidified incubator at 37 °C with 5% CO_2_ for 2–3 days. Standard and IL-coated LNPs containing Cy5-labeled siRNA were prepared as previously described and diluted in the respective growth medium to achieve a final siRNA concentration of 50 nM per well. Cells were treated with 350 µL of the prepared LNP mixtures for 4 or 24 hours. The treatment mixtures included either standard LNPs or IL-coated LNPs coated with 5, 10, or 20 µL of 1:3 or 1:5 cation: anion choline *trans*-2-hexenoate IL. After 4 or 24 hours, cells were washed with 500 µL of 1x PBS, detached using 150 µL Trypsin-EDTA solution, and collected into microcentrifuge tubes. The suspensions were centrifuged at 900 × g for 10 minutes, and the cell pellets were resuspended in flow cytometry running buffer: 500 µL of 5% FBS and 2 mM EDTA in 1*x* PBS for b.End3 cells, and 5% FBS in 1*x* PBS for NSC-34 cells. Flow cytometry analysis was conducted using the Attune NxT Acoustic Focusing Cytometer (Singapore) with Attune NxT software. Cy5 fluorescence intensity was measured with excitation at 638 nm and emission detected at 720/30 nm. A total of 30,000 events were recorded per sample. Forward *vs.* side scatter plots were used to gate live cells, excluding dead cells and debris. Untreated cells served as negative controls to account for autofluorescence and to establish gating for Cy5-negative and Cy5-positive populations. Data are expressed as the - fold increase in the percentage of Cy5-positive events, normalized to untreated controls for each treatment group.

### RBC hitchhiking study

Both whole BALB/c mouse blood and human whole blood (gender and donor-pooled with K_2_EDTA anti-coagulant, Bio-IVT, USA) were treated in triplicate at a 1:3 v/v ratio with either 1x PBS (Gibco, pH 7.4, Thermo Scientific, # 10010031), uncoated LNPs, or 1:3 IL-coated LNPs and rotary incubated at 37°C for 20 min at 50 RPM (Benchmark Scientific, #H2024). Incubated whole blood with LNPs was then spun down at 1000xg at 4°C for 10 mins in a temperature-controlled centrifuge (Thermo Scientific Sorvall Ultracentrifuge, Cat # ST8R). Subsequently, the blood components (serum, platelets, white blood cells, RBCs) were manually separated to isolate the RBC fraction, which was then washed twice as much with 1x PBS to remove unbound NPs at 200xg and 4°C for 5 mins. Washed live RBCs were then resuspended to 10% hematocrit for flow cytometric evaluation (Attune Nxt Flow Cytometer with far-red fluorescence, RL1-A) of % gated viable RBC singlets (gated from SSC-A vs. FSC-A, and validated by FSC-H *vs*. FSC-A) colocalized with one or more fluorescent NP events (side scatter SSC-A *vs*. RL1-A), which was then validated by fluorescent plate reader (excitation 640 nm/emission 670 nm, BioTek H1 Synergy Hybrid Multi-mode, Gen5 software) using a 96-well black plate (Greiner Bio-One, #655076), to calculate % of administered LNPs accumulated in the RBC fraction from the whole blood treatment dose.

### Scanning electron microscopy (SEM)

Red blood cells were initially fixed overnight in a 5 mL Eppendorf tube using a solution containing 2.5% glutaraldehyde and 2% paraformaldehyde in 1 M cacodylate buffer. Following fixation, approximately 10 µL of the cell suspension was transferred onto coverslips coated with cationic poly-L-lysine. The cells were allowed to adhere for 1 hour at 25 °C, then rinsed with 1 M cacodylate buffer. Subsequently, the samples underwent post-fixation with 2% osmium tetroxide for 1 hour at room temperature, followed by another rinse with the same buffer. Dehydration was carried out using a graded ethanol series (50%, 75%, and 100%), with drying at 25 °C after each step. After dehydration, the samples were treated with hexamethyldisilazane through multiple exchanges and allowed to dry completely at room temperature. The dried specimens were mounted on 9.5 × 9.5 mm aluminum stubs using carbon adhesive tape, with a final application of hexamethyldisilazane to ensure complete drying. To enhance imaging contrast, the samples were sputter-coated with a palladium-gold alloy (16.5 mm, 35 mA, 200 s) and stored overnight at 4 °C. Imaging was performed using a JSM-7200 FLV Field-Emission Scanning Electron Microscope.

### Hemolysis study

Whole BALB/c mouse blood and human whole blood was spun down in a centrifuge at 1000xg at 4°C for 10 mins before the blood components, other than the RBCs, were discarded. The RBCs were then washed with 1x PBS at a pH 7.4 at 500xg two more times. After the last wash, the supernatants were discarded, and RBC pellets were left. Ten μL of RBC pellet was added to 490 μL of 1x PBS at a pH of 7.4 and inverted several times to create the RBC stock solutions. RBC stock solution was added to each well of a 96 well plate according to the doses of 1:120, 1:100, 1:80, 1:40, 1:20, and 1:10 v/v in quadruplet for LNPs (uncoated and IL-coated). Twenty v/v% Triton X and 1x PBS at pH 7.4 were used as positive and negative controls, respectively, at the same dosing. After each dose was added to their respective wells, the plate was incubated at 37°C for one hour and then spun down in a centrifuge at 100xg and 4°C for 10 min. One hundred μL of supernatant of each well was then re-plated into a clean, 96 well clear plate and read on a plate reader at an absorbance of 410 nm. Hemolysis was calculated using Triton X as 100% toxic to RBCs.

### *In vivo* biodistribution and whole blood pharmacokinetics

For potential therapeutic applications, the efficacy of 1:3 IL-coated LNPs were tested by studying their biodistribution and pharmacokinetics in BALB/c mice. Four to five concentrated batches of Cy5-labeled siRNA-loaded uncoated or 1:3 IL-coated LNPs were diluted to 1.5 mL with 0.9% saline (Teknova, S5819). One hundred μL of either uncoated or 1:3 IL-coated LNP containing Cy5-labaled siRNA stocks were intravenously administered via tail vein injection into healthy BALB/c mice (n=7) and 72 h later organ accumulation and whole blood pharmacokinetics were evaluated. Live mouse imaging was taken using IVIS Lumina III (PerkinElmer) and Living Image 4.7.4 Software at five time points (3, 24, 36, 48, 72 h). The images were taken with 640/670 nm lasers at a 0.5 min exposure time and a height of 1.6 cm. In addition, global background subtraction was applied.

For whole blood pharmacokinetics, 20 μL blood draws from the submandibular vein were taken at 20 min, 1 h, 6 h, 24 h, 48 h, and 72 h using a 5 mm lancet (Braintree Scientific, INC., GR 5MM). The 20 μL whole blood was diluted to 200 μL with 0.9% saline in K_2_EDTA-coated tubes (BD, #365974) before measuring fluorescence intensities on a plate reader using a black 96-well plate (Greiner Bio-One, #655076), using 200 μL of bare and IL-coated LNPs as well as 0.9% saline as controls at 640 and 670 nm. To obtain a percent injected dose (%**ID**), the raw fluorescent readings were normalized to volume and dilution within circulating blood at each time point before being divided by injected dose fluorescence. For biodistribution, nine organs (heart, spleen, pancreas, intestines, brain, liver, kidneys, lungs, and spine) were collected, imaged on IVIS Lumina III, and stored at -80°C until homogenization. For homogenization, the organs were weighed, placed in RIPA lysis buffer (G Biosciences, #786-756) (1:1 v/v), and sonicated for 1 h in the dark. The heart, spleen, and kidney were spliced into very small pieces due to their muscular rigidity before homogenization. All tissues were emulsified using an IKA T10 Basic Ultra-turrax 5G or 10G homogenizer (Cole-Parmer, EW-04720-51) before being centrifuged at either 3260 RPM or 10,000xg at 4°C for 10 minutes depending on size of tubes. The supernatants were collected and total volumes were measured. Two hundred μL of each supernatant was pipetted into a black, 96-well plate and their fluorescence were read on a plate reader at 640 and 670 nm with saline supernatants and 200 µL LNPs as controls. The raw fluorescence was normalized to the total volume of supernatants and divided by the injected dose fluorescence to calculate % injected dose per whole organ.

## Statistical analysis

Data are presented as mean ± standard deviation (**SD**), with the number of replicates specified for each experiment. Statistical analyses were conducted using one-way or two-way ANOVA in GraphPad Prism 9 (GraphPad Software, San Diego, CA). Bonferroni’s multiple comparisons test was applied for one-way ANOVA, while Šídák’s or Tukey’s multiple comparisons tests were used for two-way ANOVA, as appropriate. The significance level (α) was set at 0.05.

## Results and discussion

### Coating LNPs with 1:3 and 1:5 choline *trans*-2-hexenoate IL resulted in significant changes in their colloidal properties

We used a bath sonication protocol that we previously developed [34] to coat siRNA-LNPs with 1:3 or 1:5 choline *trans*-2-hexenoate IL. Choline *trans*-2-hexenoate comprises a choline cation and a 2-hexenoic acid anion whereas the LNPs are positively charged with a zeta potential of +19 mV. Bath sonication and electrostatic interactions of IL with LNPs can drive the charged-based layering of ions in the IL onto positively charged LNP surfaces [24]. First, we bath sonicated LNPs for one minute to deagglomerate the particles prior to adding IL. Next, we added 5, 10, or 20 µL of 1:3 or 1:5 IL to the LNPs and bath sonicated the mixture for 60 minutes. Significant changes in the hydrodynamic diameters and/or zeta potential were used as determinants of IL-coating. A significant decrease in the sizes of IL-coated LNPs compared to uncoated LNPs might allow internalization of smaller IL-LNPs into the neural cells [35–38]. Bath sonication can deagglomerate LNPs and potentially reduce their sizes followed by IL-coating likely due to the electrostatic- and steric-based stabilizing effects provided by ILs [39]. Another influencing factor could be the formation of one or more ionic shells surrounding the LNPs, which may affect the particle size of the resulting IL-coated LNPs.

Significant variations were observed in the particle sizes of LNPs following coating with 1:3 or 1:5 IL (**Figure 2**). Uncoated LNPs exhibited an average particle diameter of 324 ± 7 nm. LNPs coated with 5 µL of 1:3 and 1:5 IL reduced the particle sizes to 184 ± 9.4 nm and 210 ± 6.4 nm, respectively (**Figure 2a**). However, increasing the coating volumes to 10 and 20 µL of 1:3 IL resulted in particle size increases to 285 ± 16.6 nm and 461 ± 46.7 nm (**Figure 2a**). Similarly, coating with 10 and 20 µL of 1:5 IL led to particle sizes of 610 ± 67 nm and 622 ± 178.3 nm, respectively. Changes in the zeta potential of coated LNPs are important to track, as they are suggestive of changes in the surface charge post-coating. The positive charge on uncoated LNPs serves as a driving force for the anionic and cationic clusters in ILs to form a coating on the nanoparticle surface. Interestingly, the addition of 5 µL of 1:3 or 1:5 IL led to a notable decrease in particle size (∼40% decrease) and dispersity indices (∼35% decrease) (**Figure 2a and b**). In contrast, the addition of 10 or 20 µL of IL resulted in a ∼50% increase in these parameters. This behavior may be attributed to the presence of excess, loosely bound ILs that engage in non-specific interactions—such as electrostatic forces or hydrogen bonding—with uncoated, partially coated, or already IL-coated LNPs. These interactions likely influence the overall colloidal properties of the LNPs, highlighting the multifaceted effects of IL coating volumes on nanoparticle behavior.

**Figure 2.**
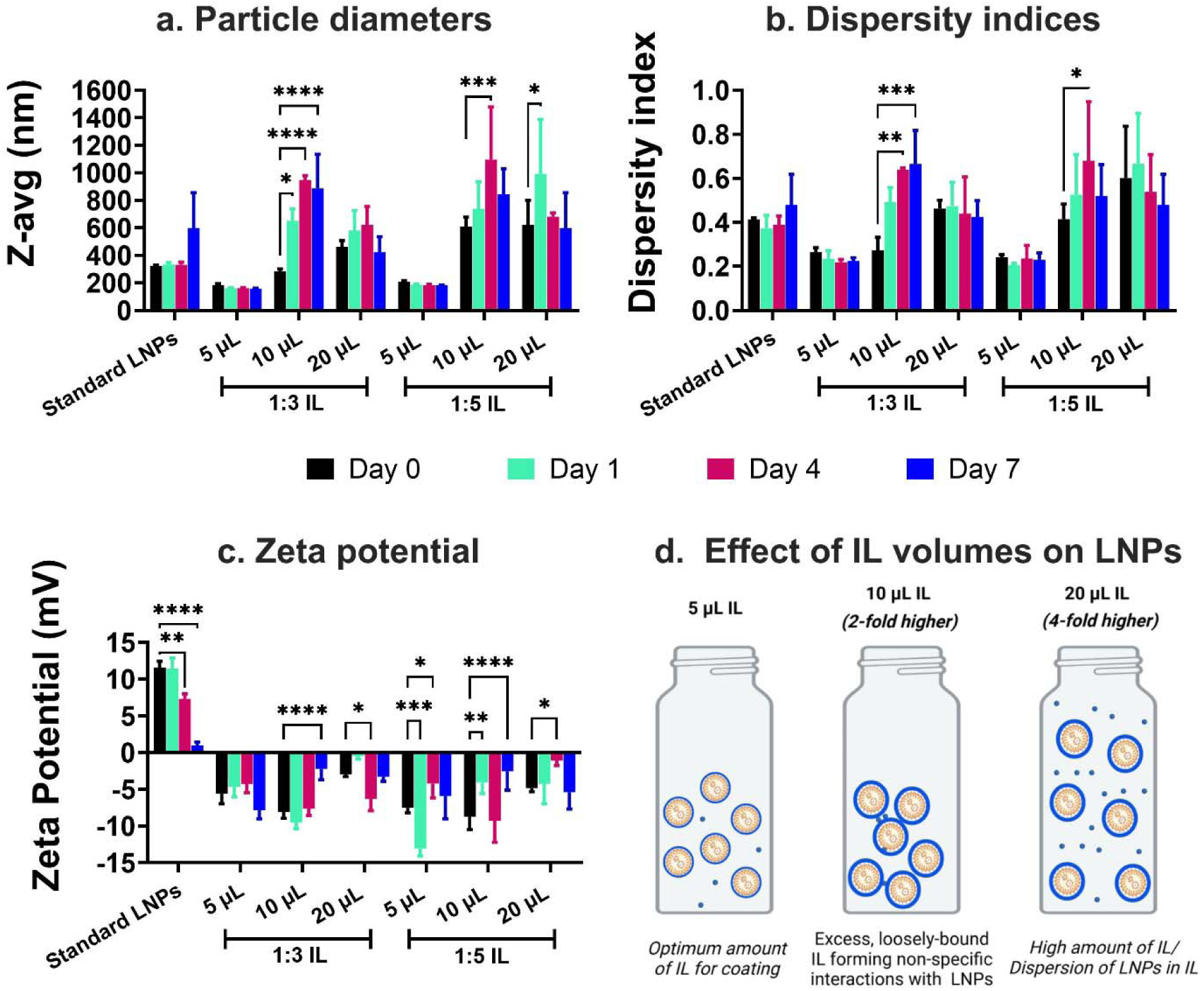
Particle diameters (a), dispersity indices (b), zeta potential (c) of standard, 1:3 and 1:5 IL-coated siRNA-LNPs over a period of seven days measured using dynamic light scattering and (d) predicted effect of different IL volumes on LNPs. siRNA-LNPs (400 nM) were initially prepared in DI water and further diluted 10*x* using nuclease-free water. 5, 10, and 20 µL of 1:3 and 1:5 choline *trans*-2-hexenoate IL were added to the siRNA-LNPs followed by bath sonication for 60 minutes. Z-average particle diameters, dispersity indices, and zeta potential were measured on days 0 (the day of preparation), 1, 4, and 7 upon storage at 2-8 °C using a Malvern Zetasizer Pro. Statistical analysis was done using two-way ANOVA with Dunnett’s multiple comparisons test. Data are presented as mean ± SD of n=3 measurements. *p<0.05, **p<0.01, ***p<0.001, and ****p<0.0001.

The introduction of charged species like ILs may further influence colloidal stability by promoting particle aggregation through electrostatic attractions between IL-coated LNPs. We also tested LNPs coated with 1:3 and 1:5 IL at different volumes (5, 10, and 20 µL) and measured changes in particle size, dispersity indices, and zeta potential over seven days with interim storage at 2–8 °C. For 1:3 IL-coated LNPs, no significant changes were observed in particle diameters for standard LNPs or those coated with 5 and 20 µL of IL (**Figure 2a**). However, LNPs coated with 10 µL IL exhibited a dramatic increase in particle size, from 285 ± 16.6 nm on day 0 (day of preparation) to 887 ± 247.6 nm on day 7. Similarly, dispersity indices for standard LNPs and LNPs coated with 5 and 20 µL of IL remained stable, while LNPs coated with 10 µL IL showed a significant increase, from 0.27 on day 0 to 0.66 on day 7 (**Figure 2b**). For zeta potential, LNPs coated with 5 µL IL exhibited no significant changes, whereas uncoated LNPs and those coated with 10 and 20 µL IL showed substantial variations over the seven-day period (**Figure 2c**). The colloidal stability of LNPs coated with 1:5 IL was similarly assessed. The excess anions in the 1:5 IL formulation could either enhance colloidal stability by increasing surface repulsion between anionic particles or promote aggregation through non-specific electrostatic and hydrogen bonding interactions. Standard LNPs and those coated with 5 and 20 µL of 1:5 IL displayed no significant changes in particle size over seven days (**Figure 2a**). However, LNPs coated with 10 µL IL showed a pronounced increase in particle size indicative of aggregation, from 610 ± 67 nm on day 0 to 1096 ± 383.1 nm on day 7 (**Figure 2a**). Dispersity indices remained stable for standard LNPs and those coated with 5 and 20 µL IL, but LNPs coated with 10 µL IL showed a notable increase, from 0.41 on day 0 to 0.68 on day 7 (**Figure 2b**). Zeta potential measurements indicated significant changes in surface charge for uncoated LNPs and those coated with 5, 10, and 20 µL IL over the storage period (**Figure 2c**). Overall, significant changes were observed in the particle sizes and dispersity indices of LNPs coated with an intermediate volume of IL (10 µL), while LNPs coated with lower (5 µL) and higher (20 µL) volumes maintained their colloidal properties over the seven-day storage period. These findings suggest that there is an optimal IL volume for coating LNPs, with 5 µL resulting in the most stable colloidal attributes. The substantial increase in particle size and dispersity indices for LNPs coated with 10 µL IL may be attributed to excess, loosely bound IL molecules creating non-specific interactions, such as electrostatic or hydrogen bonding, with uncoated, partially coated, or previously coated LNPs. Conversely, when an even larger volume of IL (20 µL) is used, the excess IL likely creates a dispersion resembling LNPs suspended within IL, potentially stabilizing the particles (**Figure 2d**). However, the larger particle size in this case may result from the formation of multiple IL layers around the LNPs. This trend was consistent with our previous study, where LNPs coated with 12.5 µL of 1:1 IL demonstrated greater colloidal stability compared to those coated with higher IL volumes [34].

Self-assembled polyelectrolyte complexes like LNPs are susceptible to aggregation and instability over time, which can compromise their therapeutic efficacy. LNPs were originally designed for efficient drug delivery to the liver, where they have shown high uptake efficiency [7, 40–45]. This hepatic targeting is primarily mediated by the adsorption of apolipoprotein E (**ApoE**) onto the LNP surface following intravenous administration [12]. The resulting ApoE-LNP complexes act as ligands for lipoprotein receptors on hepatocytes, facilitating receptor-mediated uptake and accumulation in liver cells. While this mechanism has been highly effective for liver-specific drug delivery, it limits the potential application of LNPs for targeting other tissues or organs. To overcome such limitations, surface modifications of LNPs have been explored, with ILs emerging as a promising class of materials [11, 34, 46–49]. ILs have demonstrated significant potential in enhancing drug delivery to traditionally challenging sites, such as intranasal and transdermal routes, by modulating surface interactions and improving bioavailability [19–21, 50, 51]. By leveraging their unique physicochemical properties, ILs offer an opportunity to re-engineer LNPs to alter their protein-binding profiles and redirect their biodistribution [29, 31, 52, 53]. In this context, we hypothesize that coating LNPs with biocompatible ILs could not only reduce unwanted interactions with serum proteins but also enhance their “stealth” properties. This modification may allow the LNPs to evade rapid clearance mechanisms, thereby enabling more effective delivery to distant targets such as the BBB. Specifically, IL-coated LNPs could improve accumulation in BECs lining the BBB, offering a novel approach to drug delivery for central nervous system disorders [31]. This strategy represents a significant advancement in expanding the utility of LNP technology beyond its traditional applications.

### IL-coated LNPs are cytocompatible with mouse brain endothelial and neuroblastoma cells

We evaluated the cytocompatibility of LNPs with a mouse brain endothelial cell line (bEnd.3) and mouse neuroblastoma cells (NSC-34) (**Figure 3**). For bEnd.3 cells, the viability after treatment with standard LNPs and 5, 10, and 20 µL of 1:3 IL-coated LNPs was 101.4 ± 4.9%, 92.7 ± 3.3%, 90.3 ± 8.9%, and 78.5 ± 24.5%, respectively (**Figure 3a**). When treated with 20 µL of 1:5 IL-coated LNPs, cell viability was 96.9 ± 9.6% (**Figure 3a**). Similarly, for NSC-34 cells, viability after treatment with standard LNPs and 5, 10, and 20 µL of 1:3 IL-coated LNPs was 103.6 ± 3.1%, 99.1 ± 3.1%, 93.3 ± 9.9%, and 61.7 ± 18.1%, respectively, while 20 µL of 1:5 IL-coated LNPs resulted in 99.5 ± 3.5% viability (**Figure 3b**).

**Figure 3.**
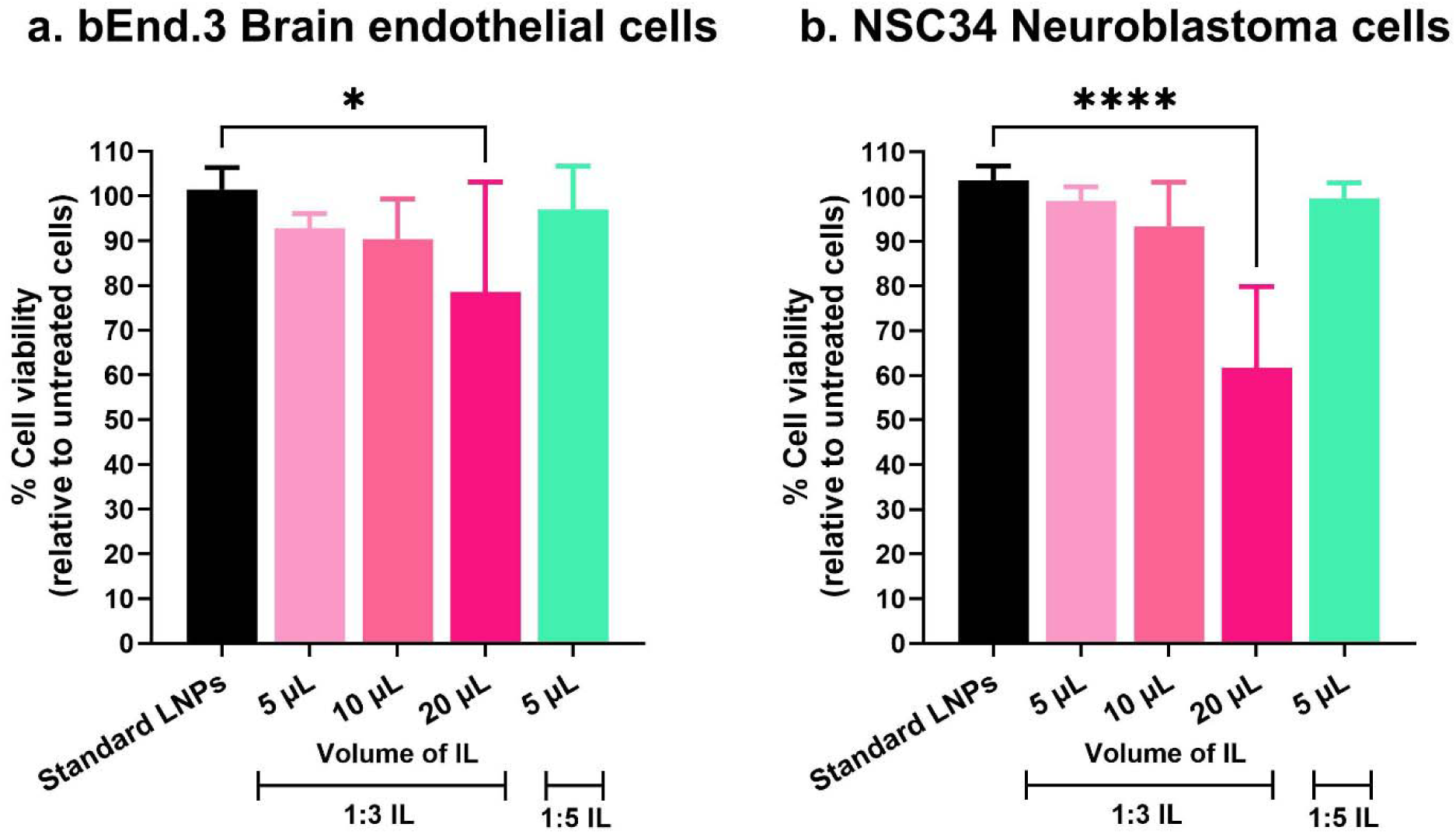
Cytocompatibility of standard and IL-coated LNPs with a mouse brain endothelial cell line, bEnd.3 (a) and NSC-34 mouse neuroblastoma cells. **(b)**. Cells were incubated with standard LNPs, 1:3 and 1:5 IL-coated LNPs containing 50 nM siGFP for four hours. Untreated cells were used as a negative control. Cell viability was determined 24 h-post-transfection using a CellGlo luminescence viability assay and the data were normalized to untreated cells. Data represents mean + SD (n=6). *p<0.05 and ****p<0.0001.

A significant reduction in cell viability was observed in both cell lines treated with 20 µL of 1:3 and 1:5 IL-coated LNPs compared to standard LNPs (*p* < 0.05 and *p* < 0.0001, respectively) (**Figure 3**). Furthermore, cell viability was inversely proportional to the volume of IL used for coating, with the greatest reduction observed for cells treated with 20 µL of 1:3 and 1:5 IL-oated LNPs. It is likely that the free/uncoated IL remaining in the suspension causes cell membrane toxicities manifesting as reduced cell viability.

### 1:3 and 1:5 IL-coated LNPs show superior uptake into mouse brain endothelial cells and mouse neuroblastoma cells compared to standard LNPs

After evaluating the cytocompatibility of LNPs, our next measured the cellular uptake of standard LNPs, and 1:3 and 1:5 IL-coated LNPs into mouse bEnd.3 BECs and mouse NSC-34 neuroblastoma cells. Using flow cytometry, a quantitative single-cell analysis technique that measures light scattering and fluorescence emissions, we analyzed the uptake of Cy5-labeled siRNA-loaded LNPs [54]. Untreated and unstained cells were included in the analysis to exclude autofluorescence. bEnd.3 cells treated with standard LNPs for 24 hours resulted in an 8.4-fold increase in Cy5 siRNA uptake (**Figure 4a**). Cells treated with 1:3 IL-coated LNPs showed a significant enhancement in uptake: 34.4-fold with 5 µL IL (p < 0.0001), 21.8-fold with 10 µL IL (p < 0.0001), and 33.4-fold with 50 µL IL (p < 0.0001), compared to standard LNPs (**Figure 4a**). Similarly, 1:5 IL-coated LNPs demonstrated a notable increase, with a 40.8-fold enhancement in uptake at 5 µL IL (p < 0.0001). In NSC-34 neuroblastoma cells, treatment with standard LNPs for four hours led to a 19.2-fold increase in Cy5 siRNA uptake (**Figure 4b**). For cells treated with 1:3 IL-coated LNPs, significant increases were observed at all IL volumes tested: 28.8-fold with 5 µL IL (p < 0.01), 28.2-fold with 10 µL IL (p < 0.01), and 50.1-fold with 50 µL IL (p < 0.0001), compared to standard LNPs (**Figure 4b**). Similarly, 1:5 IL-coated LNPs showed a substantial improvement, with a 5 -fold increase in uptake at 5 µL IL (p < 0.0001) (**Figure 4b**). These findings highlight the potential of IL-coated LNPs, particularly at optimized IL volumes, to significantly enhance the cellular uptake of siRNA in both bEnd.3 and NSC-34 cell models.

**Figure 4.**
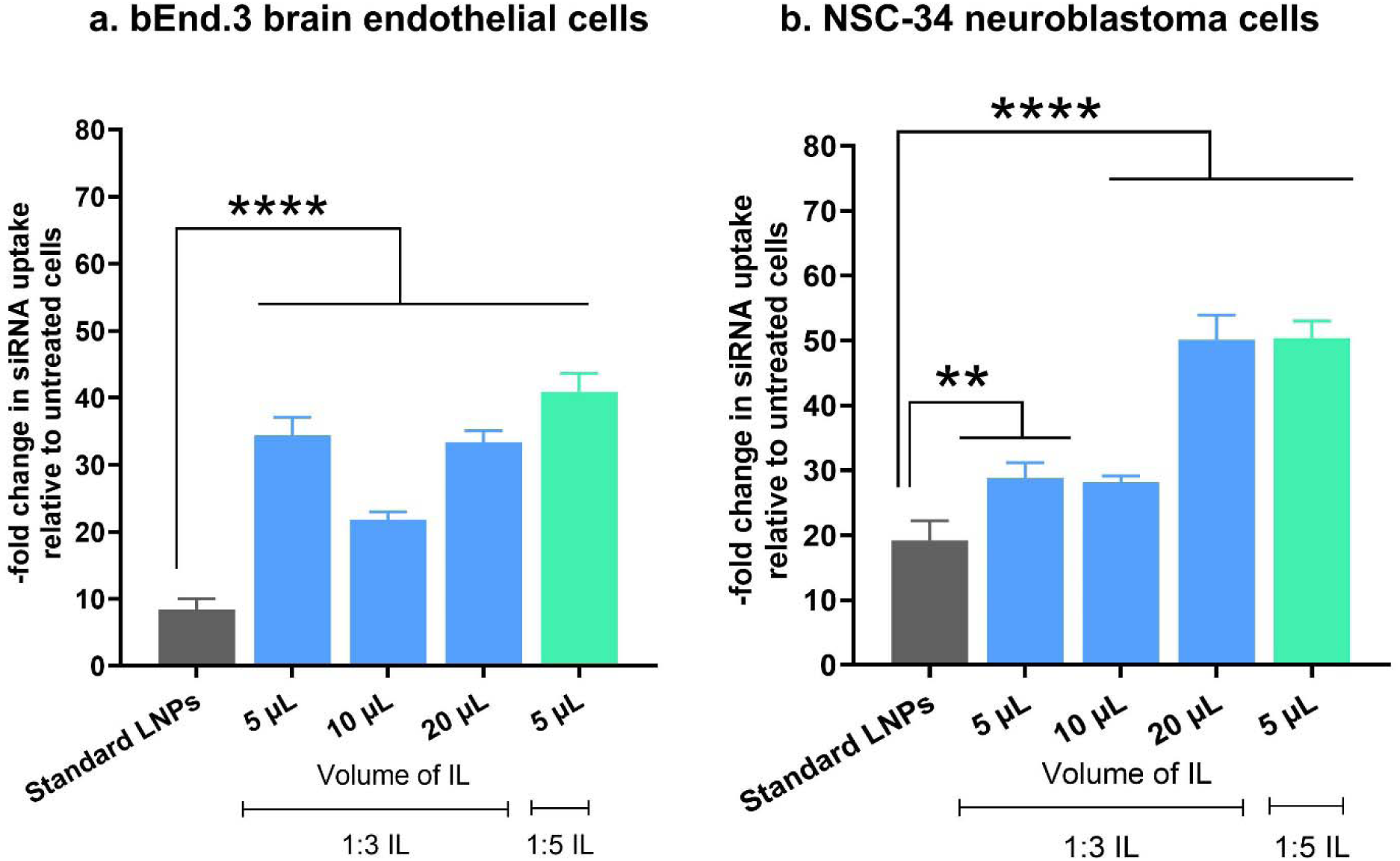
Cellular uptake of standard, 1:3 and 1:5 IL-coated LNPs containing Cy5 siRNA into bEnd.3 BECs and NSC-34 neuroblastoma cells using flow cytometry. bEnd.3 BECs and NSC-34 neuroblastoma cells were treated for 24 h and 4 h respectively with standard, 1:3 and 1:5 IL-coated LNPs loaded with 50 nM Cy5 siRNA in complete growth medium in a humidified incubator at 37 °C and 5% CO_2_. Untreated cells were used as negative controls. Data are presented as a -fold increase in Cy5 siRNA uptake (% Cy5+ cells) relative to untreated cells (n=3). Statistical analysis was done using one-way ANOVA with Dunnett’s multiple comparisons test. **p<0.01, and ****p<0.0001.

It is important to note that bEnd.3 cells used here as a model of BECs have low pinocytic capacity, necessitating extended exposure times for effective LNP uptake. In our study, we treated these cells with LNPs for 24 hours. A shorter exposure of 4 hours was also tested, but the uptake of LNPs was minimal under these conditions **(Supplementary Figure 1a)**. Notably, during the shorter treatment, standard LNPs exhibited higher uptake compared to IL-coated LNPs. This suggests that IL-coated LNPs require more time to undergo cellular uptake, as evidenced by the significantly greater uptake observed after 24 hours. In contrast, NSC-34 neuroblastoma cells, a cancer cell type, displayed the opposite trend **(Supplementary Figure 1b)**. The data presented in **Figure 4** were obtained following a 4-hour incubation with LNPs. However, when these cells were incubated for 24 hours, all LNP samples, whether standard or IL-coated, showed a substantial increase in uptake, with no apparent differences between the two. This indicates that the uptake kinetics in cancer cells are significantly faster compared to BECs, which are low pinocytic cells and exhibit slower uptake kinetics. In summary, the markedly higher uptake of Cy5 siRNA in both bEnd.3 BECs and NSC-34 neuroblastoma cells treated with 1:3 and 1:5 IL-coated LNPs, compared to standard LNPs, can be attributed to the enhanced membrane-penetrating properties conferred by the IL coating [55–57]. Given the promising behavior of 1:3 IL-coated LNPs in the studies above, we decided to focus on those LNPs as a lead candidate in the following studies.

### 1:3 IL-coated LNPs show minimal hemolysis of mouse and human RBCs

We determined the potential safety of bare LNPs and 1:3 IL-coated LNPs using mouse and human RBCs at various LNP: RBC volumetric ratios (1:120, 1:100, 1:80, 1:40, 1:20, and 1:10). As shown in **Figure 5**, we observed a dose-dependent increase in the hemolysis of RBCs. The hemolysis for mouse RBCs treated with bare LNPs were 3.3%, 0%, 2.8%, 0%, 1.2%, and 8.4% at these ratios, respectively, while those treated with 1:3 IL-coated LNPs exhibited 1.7%, 2.1%, 0.6%, 3.7%, 6.7%, and 11.8% hemolysis, respectively (**Figure 5a**). Similarly, for human RBCs, hemolysis was 0.6%, 0.2%, 0%, 0.5%, 0.2%, and 0.4% for bare LNPs and 0%, 0.5%, 0.7%, 1.6%, 4.2%, and 13.8% for 1:3 IL-coated LNPs at the corresponding volumetric ratios (**Figure 5b**).

**Figure 5.**
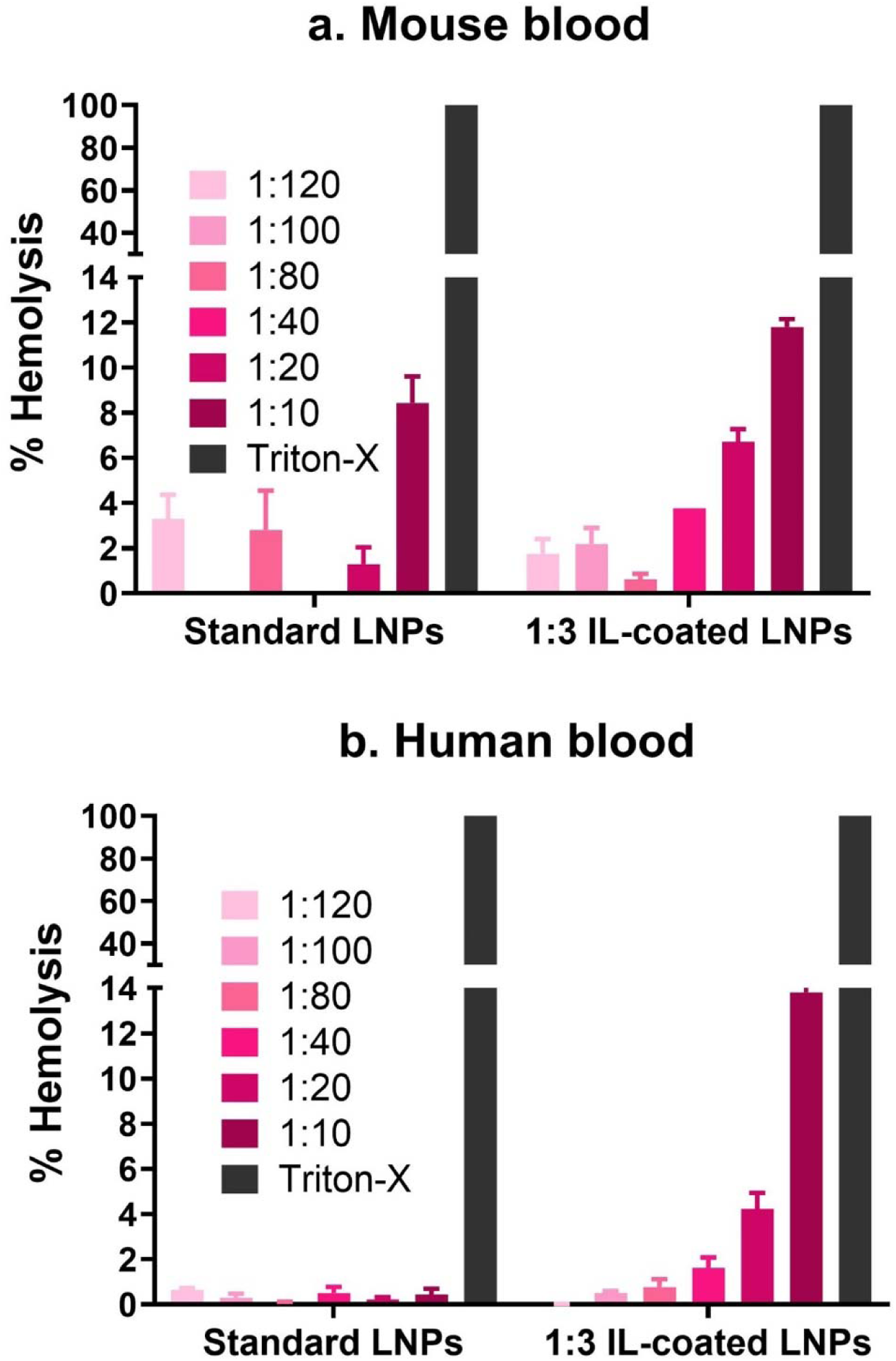
Hemolytic potential of bare or 1:3 IL-coated LNPs with mouse (a) or human (b) blood. RBC stock solution was added to a 96-well plate according to the doses of 1:120, 1:100, 1:80, 1:40, 1:20, and 1:10 v/v (LNP: RBC) for standard and 1:3 IL-coated LNPs. 20 v/v% Triton X and 1x PBS at pH 7.4 were used as positive and negative controls, respectively. Hemolysis was calculated using Triton X as 100% toxic to RBCs.

The hemolytic potential represents the likelihood of LNPs to cause RBC lysis, an undesirable outcome. Since ILs have a known affinity for RBC membranes, their potential to induce hemolysis via membrane adherence and entry is a critical factor to assess. While IL-coating enhances LNPs with RBC-hitchhiking properties, the alternating charge composition of the IL layers on the LNP surface (comprising cations and anions) is expected to reduce their ability to penetrate RBC membranes and cause hemolysis [24].

### 1:3 IL-coated LNPs show enhanced hitchhiking on mouse and human RBCs

To test whether IL-coated LNPs can attach to RBCs, we incubated Cy5-labeled siRNA-loaded LNPs with whole mouse and human blood followed by RBC pellet isolation. Cy5 fluorescence intensities of RBCs were subsequently measured using a fluorescence plate assay to calculate the percentage of total LNPs that successfully hitchhiked on RBCs. Additionally, flow cytometry was employed to quantify the percentage of RBC singlets carrying attached Cy5 siRNA-loaded LNPs (**Figure 6**, **Supplementary Figure 2**), providing further confirmation of LNP attachment to RBC surfaces.

**Figure 6.**
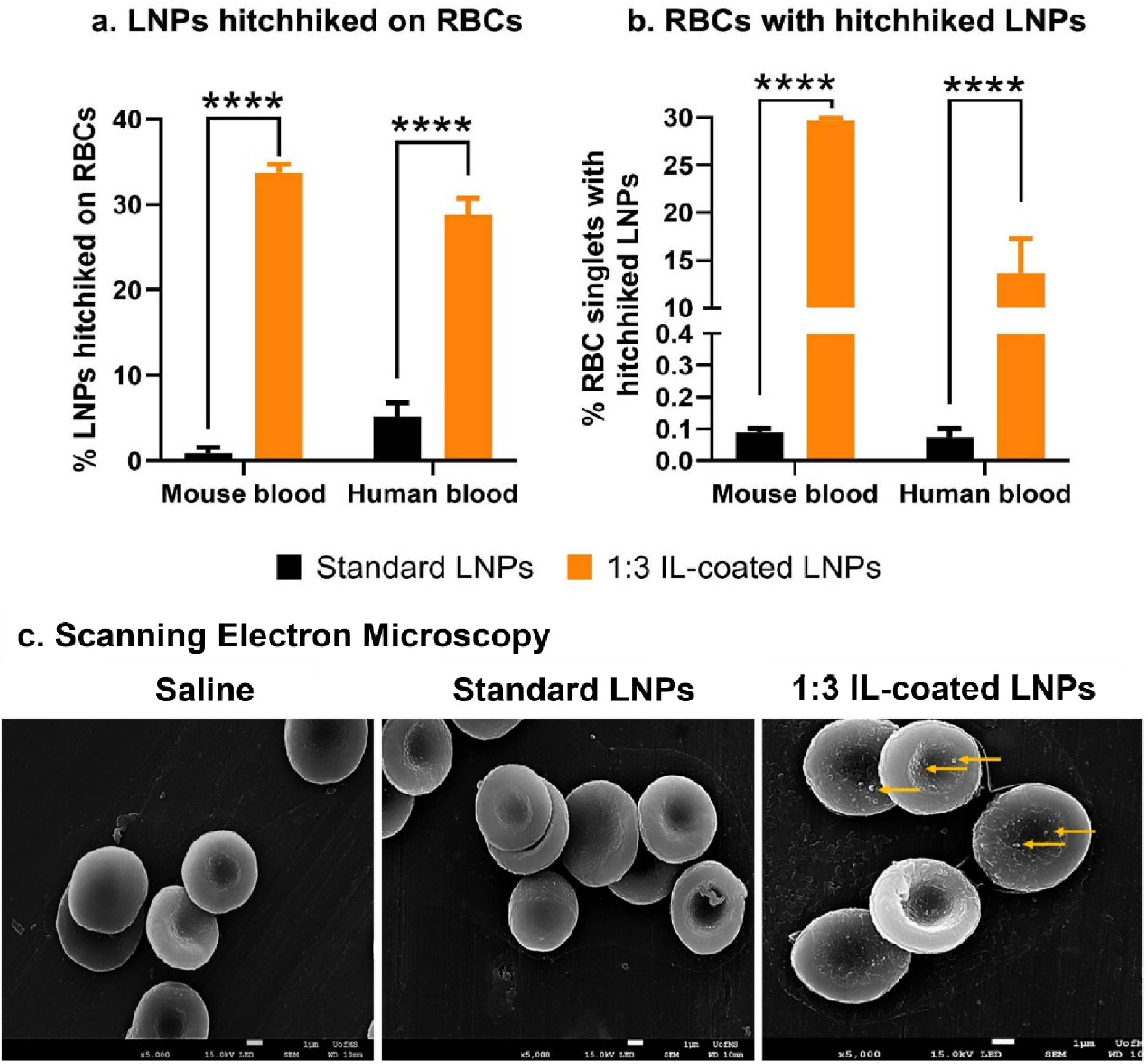
Percent of total LNPs hitchhiking on RBCs by fluorescent plate reader (a), percentage of RBC singlets with hitchhiked LNPs by flow cytometry (b) and SEM micrographs showing hitchhiked LNPs on RBCs (c) for standard (uncoated) and 1:3 IL-coated LNPs. Hitchhiking percentages were determined using (**a**) a fluorescence plate assay, where Cy5-siRNA-loaded LNPs were mixed with whole blood (1:3 v/v RBC: LNP), rotary-incubated at 37°C for 20 minutes, and RBC-associated fluorescence was measured, and (**b**) flow cytometry to assess RBC singlets with LNPs under similar conditions. RBCs were incubated with either saline, standard or IL-coated LNPs and were washed and imaged on a scanning electron microscope. IL-coated LNP attachment onto the RBC surfaces is shown with yellow arrows in panel (**c**). Data are presented as mean ± SD. Statistical significance was assessed using one-way ANOVA with Dunnett’s test (****p < 0.0001).

Using a fluorescence plate-based assay, we observed that approximately 1% of standard LNPs and ∼34% of LNPs coated with 5 µL of 1:3 IL hitchhiked on mouse RBCs (**Figure 6a**). Similarly, ∼5% of standard LNPs and ∼29% of 5 µL 1:3 IL-coated LNPs hitchhiked on human RBCs (p<0.0001) (**Figure 6a**). This represents a 34-fold increase in mouse RBC hitchhiking and a 6-fold increase in human RBC hitchhiking for 5 µL 1:3 IL-coated LNPs compared to standard LNPs. To further investigate this hitchhiking behavior, we used flow cytometry to analyze the percentage of RBC singlets carrying hitchhiked LNPs. For mouse blood, ∼0.1% of RBC singlets carried hitchhiked LNPs with standard LNP treatment, whereas ∼30% carried hitchhiked LNPs when treated with 5 µL of 1:3 IL-coated LNPs (**Figure 6b**). In human blood, ∼0.1% of RBC singlets showed hitchhiking with standard LNPs, compared to ∼14% with 5 µL 1:3 IL-coated LNPs (significant difference, p<0.0001) (**Figure 6b**). These results correspond to a 300-fold increase in mouse RBC singlet hitchhiking and a 140-fold increase in human RBC singlet hitchhiking with 5 µL 1:3 IL-coated LNPs compared to standard LNPs. Additionally, scanning electron microscopy analysis was also performed to confirm hitchhiking of IL-coated LNPs on RBCs. As shown in **Figure 6c**, IL-coated LNPs were visibly associated with the surfaces of RBCs, whereas RBCs treated with standard LNPs appeared free of nanoparticle attachment, similar to those treated with control, saline.

The RBC hitchhiking potential of IL-coated LNPs is derived from the affinity of the 2-hexenoic acid anion component of the IL for the monocarboxylate transporter-1 in the RBC membrane [58]. The potential of IL-coating to redirect LNP biodistribution to distant sites such as the BBB is contingent on the RBC-hitchhiking ability of IL-coated LNPs. If the IL-coated LNPs undergo RBC-hitchhiking, a comparatively higher dose can likely reach the BBB along with the RBCs [59]. Previous studies have reported the RBC hitchhiking abilities of polymeric PLGA nanoparticles coated with choline *trans*-2-hexenoate IL [25, 26]. Similarly, we tested whether a coating of IL can confer RBC hitchhiking properties to LNPs. Thus, we studied the *ex vivo* RBC-hitchhiking potential of IL-coated LNPs with whole mouse and human blood to determine whether RBCs can act as “taxis” to shuttle membrane-adsorbed LNPs to the BBB. Although IL-coating confers LNPs with RBC-hitchhiking properties, it should be noted that the alternating charge composition (alternating cation and anion layers) of IL on the LNP surface will potentially evade their entry into the RBCs [24]. This implies that while IL-coated LNPs can attach to RBC membranes, they have a lesser propensity to enter the RBCs. The mild permeabilization ability of IL in addition to the affinity of 2-hexenoic acid to the RBC membrane can potentially drive the attachment of IL-coated LNPs on RBCs enabling RBC-hitchhiking.

### IL-coating prolongs blood retention, lowers liver uptake and routes LNPs to the brain

We next investigated whether enhanced RBC hitchhiking of IL-coated LNPs could prolong blood circulation and improve siRNA routing to the brain. Our previous studies demonstrated that IL-coating reduces plasma protein adsorption onto LNP surfaces—a key factor contributing to hepatic uptake [11, 34]. We hypothesized that reduced protein adsorption would result in lower liver accumulation and extended systemic circulation, thereby facilitating delivery to distant organs such as the brain. To test this hypothesis, we performed *in vivo* whole blood pharmacokinetic and biodistribution studies in BALB/c mice. Mice were intravenously injected with Cy5-labeled siRNA encapsulated in either standard or 1:3 IL-coated LNPs via the tail vein. Blood samples were collected at 0.5, 1, 6, 24, 48, and 72 hours post-injection via submandibular vein. Fluorescence intensity was measured after dilution with 0.9% saline, and the percentage of injected dose was calculated by normalizing to blood volume and dilution, and then those values were further divided by the injected dose fluorescence intensity. As shown in **Figure 7a**, both formulations exhibited peak blood levels at 0.5 h post-injection, with standard LNPs showing 74% of the injected dose and IL-coated LNPs showing 61%. However, standard LNPs were cleared more rapidly, with blood levels dropping to 28% at 1 h and 11% at 48 h. In contrast, IL-coated LNPs maintained higher blood levels over time, with 48% at 1 h and 26% at 48 h. Area under the curve (**AUC**) analysis further confirmed this trend, with IL-coated LNPs exhibiting a significantly higher AUC (1975) compared to standard LNPs (1030), indicating greater overall exposure. These results suggest that IL-coating prolongs LNP circulation, likely due to enhanced RBC-hitchhiking and reduced plasma protein adsorption. The slower clearance of IL-coated LNPs may be attributed to their reduced interaction with plasma proteins, which are known to facilitate hepatic uptake. This extended circulation time may likely enhance delivery to distant organs such as the brain.

**Figure 7.**
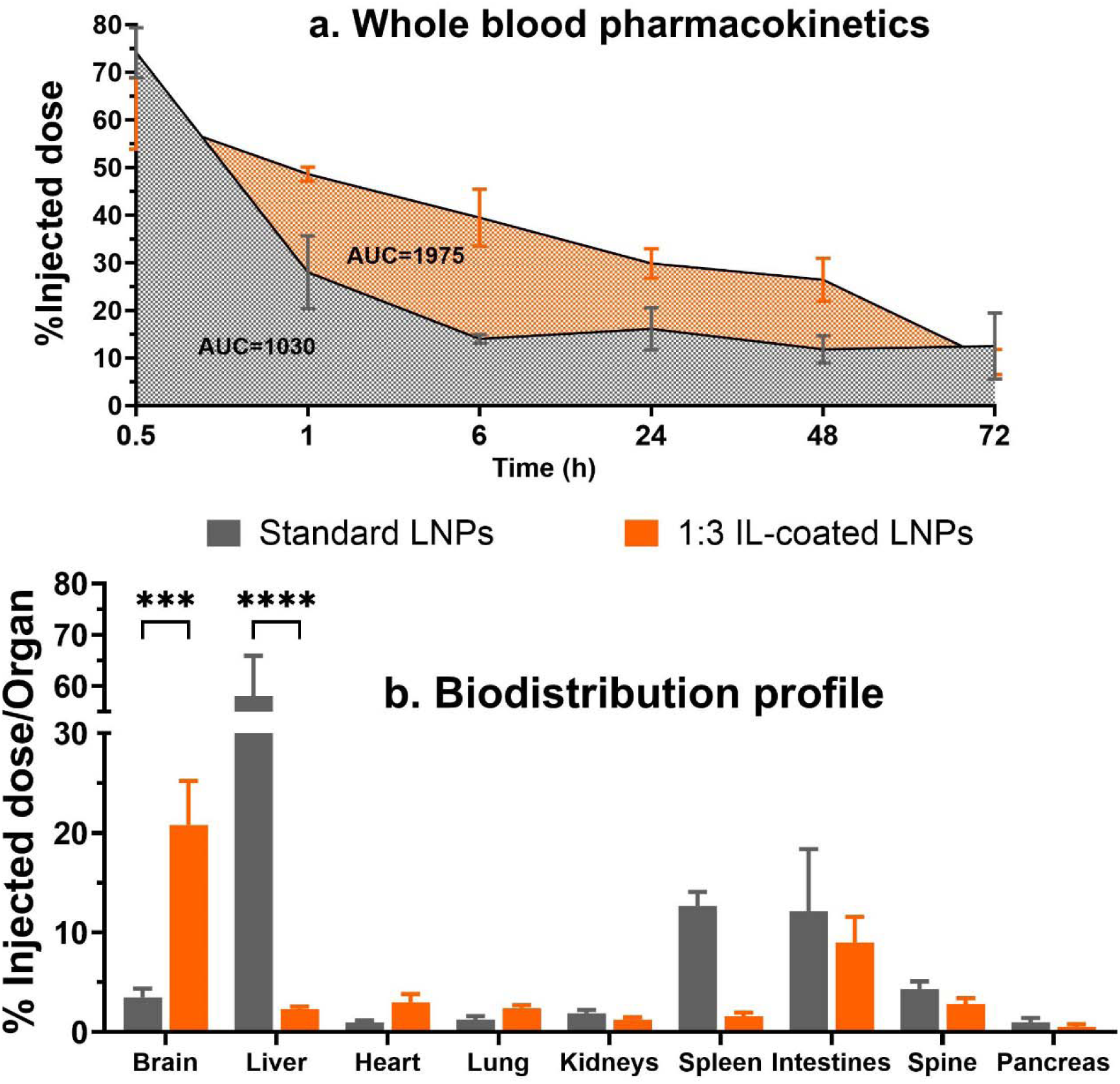
Whole blood pharmacokinetics (a) and biodistribution profile (b) of standard and 1:3 IL-coated LNPs. Standard and 1:3 IL-coated LNPs containing Cy5-labeled siRNA were injected via tail vein in mice and the % injected dose at various timepoints was calculated to determine the whole blood pharmacokinetic profile of LNPs. Additionally, the % injected dose in each organ was calculated to determine the biodistribution profile of LNPs. Data are presented as mean ± SEM. Statistical significance was assessed using two-way ANOVA with Sidak’s multiple comparisons test (***p<0.001 and ****p < 0.0001).

To assess organ-specific accumulation, major organs—including brain, liver, spleen, heart, lungs, kidneys, pancreas, intestines, and spine—were harvested 72 h post-injection. Organs were imaged using IVIS Lumina, homogenized, and analyzed for fluorescence levels. The % injected dose per organ was calculated by normalizing fluorescence to homogenate volume and injected dose. As shown in **Figure 7b**, IL-coated LNPs exhibited significantly reduced liver accumulation (2%) compared to standard LNPs (58%), consistent with reduced plasma protein adsorption and hepatic clearance. Importantly, IL-coated LNPs showed markedly enhanced brain accumulation (21%) relative to standard LNPs (3% ID), indicating successful *in vivo* translation of RBC-hitchhiking and improved routing to the brain. Given the substantial cardiac output directed to the brain, RBC-bound LNPs may preferentially route to cerebral vasculature. Additional accumulation was observed in the spleen and intestines for standard LNPs, and in the intestines for IL-coated LNPs, suggesting formulation-dependent biodistribution patterns beyond the liver and brain. These findings demonstrate that IL-coating enhances LNP circulation and brain routing while reducing hepatic uptake. The dual mechanisms of reduced plasma protein adsorption and RBC-hitchhiking contribute to the favorable pharmacokinetic and biodistribution profile of IL-coated LNPs, supporting their potential for siRNA delivery to the brain.

In our future studies, we will validate findings from the biodistribution and pharmacokinetic studies using orthogonal labeling methods such as radiolabeled IL-LNPs. Radiolabeled LNPs will allow distinguishing LNP uptake into the brain capillaries *vs*. parenchyma and determining the rates of LNP influx/efflux across the BBB [60, 61]. Followed by that, we will test the therapeutic efficacy of siRNA-loaded IL-LNPs in a mouse model of transient ischemic stroke [62, 63].

## Conclusions

We have re-engineered LNPs using 1:3 and 1:5 cation: anion ILs to develop IL-coated LNPs for siRNA delivery to brain endothelial cells and neuroblastoma cells. By optimizing the IL-coating protocol, we demonstrated that IL coating alters the colloidal properties of LNPs. LNPs coated with 5 μL of 1:3 or 1:5 IL maintained their size and dispersity indices for up to 7 days when stored at 2–8 °C. The IL-coated LNPs exhibited enhanced uptake in mouse BECs and neuroblastoma cells compared to uncoated LNPs. IL-coated LNPs achieved significantly higher levels of hitchhiking on RBCs and increased the percentage of RBC singlets involved in hitchhiking compared to standard LNPs. IL-coating of LNPs significantly enhanced systemic circulation and brain routing of siRNA by promoting RBC-hitchhiking and reducing plasma protein adsorption. This dual mechanism results in lower hepatic accumulation and improved biodistribution, highlighting the potential of IL-coated LNPs for delivery of brain therapeutics. These findings highlight the potential of IL-based re-engineering to create novel LNP carriers for delivering siRNA to challenging distant targets, such as the brain.

## Conflicts of interest

E.E.L.T. is an inventor on patents covering ionic liquids and E.E.L.T, P.K. and D.S.M are listed inventors on a patent application involving ionic liquid-modified LNPs.

## Acknowledgments.

This work was supported via discretionary research funds from the PI Manickam’s laboratory. EELT acknowledges the NSF (2236629).

## Abbreviations to ease manuscript reading

(*non-standard abbreviations used in this manuscript)

BECs: Brain endothelial cells
ILs: Ionic liquids
LNPs: Lipid nanoparticles
RBC-hitchhiking: Red blood cell hitchhiking
siRNA: Small interfering RNA
\**IL-coated LNPs*: standard LNPs coated with ILs

## Notes

### Summary of Updates

New in vivo biodistribution and pharmacokinetic data have been added

## References

1. Akinc, A., M.A. Maier, M. Manoharan, K. Fitzgerald, M. Jayaraman, S. Barros, S. Ansell, X. Du, M.J. Hope, and T.D. Madden, The Onpattro story and the clinical translation of nanomedicines containing nucleic acid-based drugs. Nature nanotechnology, 2019. 14(12): p. 1084–1087.

2. Cross, R., Without these lipid shells, there would be no mRNA vaccines for COVID-19. Chem Eng News, 2021. 99(8): p. 144.

3. Whitehead, K.A., J.R. Dorkin, A.J. Vegas, P.H. Chang, O. Veiseh, J. Matthews, O.S. Fenton, Y. Zhang, K.T. Olejnik, and V. Yesilyurt, Degradable lipid nanoparticles with predictable in vivo siRNA delivery activity. Nature communications, 2014. 5(1): p. 4277.

4. Tanne, J.H., Covid-19: FDA approves Pfizer-BioNTech vaccine in record time. 2021, British Medical Journal Publishing Group.

5. Belliveau, N., J. Huft, P. Lin, S. Chen, A. Leung, T. Leaver, A. Wild, J. Lee, R. Taylor, Y. Tam, C. Hansen, and P. Cullis, Microfluidic Synthesis of Highly Potent Limit-size Lipid Nanoparticles for In Vivo Delivery of siRNA. Molecular therapy. Nucleic acids, 2012. 1: p. e37.

6. Hou, X., T. Zaks, R. Langer, and Y. Dong, Lipid nanoparticles for mRNA delivery. Nature Reviews Materials, 2021. 6.

7. Leung, A., I. Hafez, S. Baoukina, N. Belliveau, I. Zhigaltsev, E. Afshinmanesh, D. Tieleman, C. Hansen, M. Hope, and P. Cullis, Lipid Nanoparticles Containing siRNA Synthesized by Microfluidic Mixing Exhibit an Electron-Dense Nanostructured Core. The journal of physical chemistry. C, Nanomaterials and interfaces, 2012. 116: p. 18440–18450.

8. Johnsen, K.B., A. Burkhart, F. Melander, P.J. Kempen, J.B. Vejlebo, P. Siupka, M.S. Nielsen, T.L. Andresen, and T. Moos, Targeting transferrin receptors at the blood-brain barrier improves the uptake of immunoliposomes and subsequent cargo transport into the brain parenchyma. Sci Rep, 2017. 7(1): p. 10396.

9. D’Souza, A., K.M. Dave, R.A. Stetler, and D. S. Manickam, Targeting the blood-brain barrier for the delivery of stroke therapies. Advanced Drug Delivery Reviews, 2021. 171: p. 332–351.

10. Dave, K.M. S.D. B., and D.S. and Manickam, Delivery of mitochondria-containing extracellular vesicles to the BBB for ischemic stroke therapy. Expert Opinion on Drug Delivery, 2023. 20(12): p. 1769–1788.

11. Khare, P., S.X. Edgecomb, C.M. Hamadani, E.E.L. Tanner, and D. S Manickam, Lipid nanoparticle-mediated drug delivery to the brain. Advanced Drug Delivery Reviews, 2023. 197: p. 114861.

12. Mahley, R.W., Apolipoprotein E: cholesterol transport protein with expanding role in cell biology. Science, 1988. 240(4852): p. 622–630.

13. Akinc, A., W. Querbes, S. De, J. Qin, M. Frank-Kamenetsky, K. Jayaprakash, M. Jayaraman, K. Rajeev, W. Cantley, R. Dorkin, J. Butler, L. Qin, T. Racie, A. Sprague, F. Eugenio, A. Zeigerer, M. Hope, M. Zerial, D. Sah, and M. Maier, Targeted Delivery of RNAi Therapeutics With Endogenous and Exogenous Ligand-Based Mechanisms. Molecular therapy : the journal of the American Society of Gene Therapy, 2010. 18: p. 1357–64.

14. Shi, B., E. Keough, A. Matter, K. Leander, S. Young, E. Carlini, A.B. Sachs, W. Tao, M. Abrams, and B. Howell, Biodistribution of small interfering RNA at the organ and cellular levels after lipid nanoparticle-mediated delivery. Journal of Histochemistry & Cytochemistry, 2011. 59(8): p. 727–740.

15. Gebbie, M.A., A.M. Smith, H.A. Dobbs, G.G. Warr, X. Banquy, M. Valtiner, M.W. Rutland, J.N. Israelachvili, S. Perkin, and R. Atkin, Long range electrostatic forces in ionic liquids. Chemical communications, 2017. 53(7): p. 1214–1224.

16. Rudzinski, J.F., S. Kloth, S. Wörner, T. Pal, K. Kremer, T. Bereau, and M. Vogel, Dynamical properties across different coarse-grained models for ionic liquids. Journal of Physics: Condensed Matter, 2021. 33(22): p. 224001.

17. Wang, Y.-L., B. Li, S. Sarman, F. Mocci, Z.-Y. Lu, J. Yuan, A. Laaksonen, and M.D. Fayer, Microstructural and dynamical heterogeneities in ionic liquids. Chemical reviews, 2020. 120(13): p. 5798–5877.

18. Curreri, A.M., S. Mitragotri, and E.E. Tanner, Recent advances in ionic liquids in biomedicine. Advanced Science, 2021. 8(17): p. 2004819.

19. Hattori, T., H. Tagawa, M. Inai, T. Kan, S.-i. Kimura, S. Itai, S. Mitragotri, and Y. Iwao, Transdermal delivery of nobiletin using ionic liquids. Scientific Reports, 2019. 9(1): p. 1–11.

20. Tanigawa, H., N. Suzuki, and T. Suzuki, Application of ionic liquid to enhance the nose-to-brain delivery of etodolac. European Journal of Pharmaceutical Sciences, 2022. 178: p. 106290.

21. Tanner, E.E., K.N. Ibsen, and S. Mitragotri, Transdermal insulin delivery using choline-based ionic liquids (CAGE). Journal of controlled release, 2018. 286: p. 137–144.

22. Kim, J., Y. Gao, Z. Zhao, D. Rodrigues, E.E. Tanner, K. Ibsen, P.K. Sasmal, R. Jaladi, S. Alikunju, and S. Mitragotri, A deep eutectic-based, self-emulsifying subcutaneous depot system for apomorphine therapy in Parkinson’s disease. Proceedings of the National Academy of Sciences, 2022. 119(9): p. e2110450119.

23. Pang, J., Y. Luan, F. Li, X. Cai, and Z. Li, Ionic liquid-assisted synthesis of silica particles and their application in drug release. Materials letters, 2010. 64(22): p. 2509–2512.

24. Hamadani, C.M., M.J. Goetz, S. Mitragotri, and E.E. Tanner, Protein-avoidant ionic liquid (PAIL)–coated nanoparticles to increase bloodstream circulation and drive biodistribution. Science advances, 2020. 6(48): p. eabd7563.

25. Brenner, J., S. Mitragotri, and V. Muzykantov, Red Blood Cell Hitchhiking: A Novel Approach for Vascular Delivery of Nanocarriers. Annual Review of Biomedical Engineering, 2021. 23.

26. Zhao, Z., J. Kim, V.V. Chandran Suja, N. Kapate, Y. Gao, J. Guo, V. Muzykantov, and S. Mitragotri, Red Blood Cell Anchoring Enables Targeted Transduction and Re Administration of AAV Mediated Gene Therapy. Advanced Science, 2022. 9.

27. Slowing, I.I., C.W. Wu, J.L. Vivero Escoto, and V.S.Y. Lin, Mesoporous silica nanoparticles for reducing hemolytic activity towards mammalian red blood cells. Small, 2009. 5(1): p. 57–62.

28. Tanner, E.E., Ionic liquids charge ahead. Nature Chemistry, 2022. 14(7): p. 842–842.

29. Hamadani, C.M., I. Chandrasiri, M.L. Yaddehige, G.S. Dasanayake, I. Owolabi, A. Flynt, M. Hossain, L. Liberman, T.P. Lodge, T.A. Werfel, D.L. Watkins, and E.E.L. Tanner, Improved nanoformulation and bio-functionalization of linear-dendritic block copolymers with biocompatible ionic liquids. Nanoscale, 2022. 14(16): p. 6021–6036.

30. Hamadani, C.M., G.S. Dasanayake, M.E. Gorniak, M.C. Pride, W. Monroe, C.M. Chism, R. Heintz, E. Jarrett, G. Singh, S.X. Edgecomb, and E.E.L. Tanner, Development of ionic liquid-coated PLGA nanoparticles for applications in intravenous drug delivery. Nature Protocols, 2023. 18(8): p. 2509–2557.

31. Hamadani, C.M., F. Mahdi, A. Merrell, J. Flanders, R. Cao, P. Vashisth, G.S. Dasanayake, D.S. Darlington, G. Singh, and M.C. Pride, Ionic Liquid Coating Driven Nanoparticle Delivery to the Brain: Applications for NeuroHIV. Advanced Science, 2024. 11(23): p. 2305484.

32. Khare, P., J.F. Conway, and D.S. Manickam, Lipidoid nanoparticles increase ATP uptake into hypoxic brain endothelial cells. European Journal of Pharmaceutics and Biopharmaceutics, 2022. 180: p. 238–250.

33. Khare, P., K.M. Dave, Y.S. Kamte, M.A. Manoharan, L.A. O’Donnell, and D.S. Manickam, Development of lipidoid nanoparticles for siRNA delivery to neural cells. The AAPS journal, 2021. 24(1): p. 8.

34. Khare, P., S.X. Edgecomb, C.M. Hamadani, J.F. Conway, E.E.L. Tanner, and S.M. D, Ionic liquid-coated lipid nanoparticles increase siRNA uptake into CNS targets. Nanoscale Adv, 2024. 6(7): p. 1853–1873.

35. Cho, E.C., L. Au, Q. Zhang, and Y. Xia, The effects of size, shape, and surface functional group of gold nanostructures on their adsorption and internalization by cells. Small, 2010. 6(4): p. 517–522.

36. Freese, C., M.I. Gibson, H.-A. Klok, R.E. Unger, and C.J. Kirkpatrick, Size-and coating-dependent uptake of polymer-coated gold nanoparticles in primary human dermal microvascular endothelial cells. Biomacromolecules, 2012. 13(5): p. 1533–1543.

37. He, C., Y. Hu, L. Yin, C. Tang, and C. Yin, Effects of particle size and surface charge on cellular uptake and biodistribution of polymeric nanoparticles. Biomaterials, 2010. 31(13): p. 3657–3666.

38. He, Q., Z. Zhang, Y. Gao, J. Shi, and Y. Li, Intracellular localization and cytotoxicity of spherical mesoporous silica nano and microparticles. Small, 2009. 5(23): p. 2722–2729.

39. He, Z. and P. Alexandridis, Nanoparticles in ionic liquids: interactions and organization. Physical Chemistry Chemical Physics, 2015. 17(28): p. 18238–18261.

40. Akinc, A., M. Maier, M. Manoharan, K. Fitzgerald, M. Jayaraman, S. Barros, S. Ansell, X. Du, M. Hope, T. Madden, B. Mui, S. Semple, Y. Tam, M. Ciufolini, D. Witzigmann, J. Kulkarni, R. van der Meel, and P. Cullis, The Onpattro story and the clinical translation of nanomedicines containing nucleic acid-based drugs. Nature Nanotechnology, 2019. 14: p. 1084–1087.

41. Jayaraman, M., S.M. Ansell, B.L. Mui, Y.K. Tam, J. Chen, X. Du, D. Butler, L. Eltepu, S. Matsuda, and J.K. Narayanannair, Maximizing the potency of siRNA lipid nanoparticles for hepatic gene silencing in vivo. Angewandte Chemie, 2012. 124(34): p. 8657–8661.

42. Kulkarni, J., P. Cullis, and R. van der Meel, Lipid Nanoparticles Enabling Gene Therapies: From Concepts to Clinical Utility. Nucleic Acid Therapeutics, 2018. 28.

43. Love, K.T., K.P. Mahon, C.G. Levins, K.A. Whitehead, W. Querbes, J.R. Dorkin, J. Qin, W. Cantley, L.L. Qin, and T. Racie, Lipid-like materials for low-dose, in vivo gene silencing. Proceedings of the National Academy of Sciences, 2010. 107(5): p. 1864–1869.

44. Pharmaceuticals, A. Onpattro (patisiran) lipid complex injection. 2022; Available from: https://www.onpattro.com.

45. Semple, S.C., A. Akinc, J. Chen, A.P. Sandhu, B.L. Mui, C.K. Cho, D.W. Sah, D. Stebbing, E.J. Crosley, and E. Yaworski, Rational design of cationic lipids for siRNA delivery. Nature biotechnology, 2010. 28(2): p. 172–176.

46. Hamadani, C., M. Goetz, S. Mitragotri, and E. Tanner, Protein-avoidant ionic liquid (PAIL)–coated nanoparticles to increase bloodstream circulation and drive biodistribution. Science Advances, 2020. 6: p. eabd7563.

47. He, Z. and P. Alexandridis, Ionic liquid and nanoparticle hybrid systems: Emerging applications. Advances in Colloid and Interface Science, 2017. 244: p. 54–70.

48. He, Z., L. Miao, R. Jordan, S.M. D. R. Luxenhofer, and A.V. Kabanov, A Low Protein Binding Cationic Poly(2-oxazoline) as Non-Viral Vector. Macromol Biosci, 2015. 15(7): p. 1004–20.

49. Obliosca, J., I.H. Arellano, M. Huang, and S. Arco, Double layer micellar stabilization of gold nanocrystals by greener ionic liquid 1-butyl-3-methylimidazolium lauryl sulfate. Materials Letters, 2010. 64: p. 1109–1112.

50. Banerjee, A., K. Ibsen, Y. Iwao, M. Zakrewsky, and S. Mitragotri, Transdermal protein delivery using choline and geranate (CAGE) deep eutectic solvent. Advanced healthcare materials, 2017. 6(15): p. 1601411.

51. Zakrewsky, M., K.S. Lovejoy, T.L. Kern, T.E. Miller, V. Le, A. Nagy, A.M. Goumas, R.S. Iyer, R.E. Del Sesto, and A.T. Koppisch, Ionic liquids as a class of materials for transdermal delivery and pathogen neutralization. Proceedings of the National Academy of Sciences, 2014. 111(37): p. 13313–13318.

52. Hamadani, C.M., A Novel Chemotherapeutic Nanoparticle Drug Delivery System: Surface Functionalization of Polymeric Nanoparticles With Protein-Phobic Ionic Liquid to Enhance Drug Bioavailability in Systemic Circulation. 2020, Harvard University

53. Tanner, E.E., A.M. Curreri, J.P. Balkaran, N.C. Selig Wober, A.B. Yang, C. Kendig, M.P. Fluhr, N. Kim, and S. Mitragotri, Design principles of ionic liquids for transdermal drug delivery. Advanced materials, 2019. 31(27): p. 1901103.

54. Davey, H.M. and D.B. Kell, Flow cytometry and cell sorting of heterogeneous microbial populations: The importance of single-cell analyses. Microbiological Reviews, 1996. 60: p. 641–696.

55. Cook, K., K. Tarnawsky, A.J. Swinton, D.D. Yang, A.S. Senetra, G.A. Caputo, B.R. Carone, and T.D. Vaden, Correlating lipid membrane permeabilities of imidazolium ionic liquids with their cytotoxicities on yeast, bacterial, and mammalian cells. Biomolecules, 2019. 9(6): p. 251.

56. Gal, N., D. Malferarri, S. Kolusheva, P. Galletti, E. Tagliavini, and R. Jelinek, Membrane interactions of ionic liquids: Possible determinants for biological activity and toxicity. Biochimica et Biophysica Acta (BBA) - Biomembranes, 2012. 1818(12): p. 2967–2974.

57. Kaur, N., S. Kumar, Shiksha, G.K. Gahlay, and V.S. Mithu, Cytotoxicity and Membrane Permeability of Double-Chained 1,3-Dialkylimidazolium Cations in Ionic Liquids. The Journal of Physical Chemistry B, 2021. 125(14): p. 3613–3621.

58. Hamadani, C.M., G.R. Taylor, A. Cecil, C.M. Chism, E. Huff, G. Dasanayake, J. Everett, A. Kaur, G.W. Monroe, M. Patel, E. Pierpaoli, and E.E.L. Tanner, Insights into the physicochemical interactions of ionic liquid-coated polymeric nanoparticles with red blood cells. Nanoscale Adv, 2025. 7(17): p. 5273–5283.

59. Carter, R., The human brain book: An illustrated guide to its structure, function, and disorders. 2019: Penguin.

60. Banks, W.A., P. Sharma, K.M. Bullock, K.M. Hansen, N. Ludwig, and T.L. Whiteside, Transport of Extracellular Vesicles across the Blood-Brain Barrier: Brain Pharmacokinetics and Effects of Inflammation. International journal of molecular sciences, 2020. 21(12): p. 4407.

61. Klyachko, N.L., D.S. Manickam, A.M. Brynskikh, S.V. Uglanova, S. Li, S.M. Higginbotham, T.K. Bronich, E.V. Batrakova, and A.V. Kabanov, Cross-linked antioxidant nanozymes for improved delivery to CNS. Nanomedicine: Nanotechnology, Biology and Medicine, 2012. 8(1): p. 119–129 (*equal contribution).

62. Dave, K.M., D.B. Stolz, V.R. Venna, V.A. Quaicoe, M.E. Maniskas, M.J. Reynolds, R. Babidhan, D.X. Dobbins, M.N. Farinelli, A. Sullivan, T.N. Bhatia, H. Yankello, R. Reddy, Y. Bae, R.K. Leak, S.S. Shiva, L.D. McCullough, and D.S. Manickam, Mitochondria-containing extracellular vesicles (EV) reduce mouse brain infarct sizes and EV/HSP27 protect ischemic brain endothelial cultures. Journal of Controlled Release, 2023. 354: p. 368–393

63. Dave, K.M., V.R. Venna, K.S. Rao, D.B. Stolz, B. Brady, V.A. Quaicoe, M.E. Maniskas, E.E. Hildebrand, D. Green, M. Chen, J. Milosevic, S.-y. Zheng, S.S. Shiva, L.D. McCullough, and D. S Manickam, Mitochondria-containing extracellular vesicles from mouse vs. human brain endothelial cells for ischemic stroke therapy. Journal of Controlled Release, 2024. 373: p. 803–822.

