## Supplemental File for "Ionic liquid-coated lipid nanoparticles demonstrate prolonged circulation and brain uptake via red blood cell hitchhiking"

.

***Corresponding author:**	Devika S Manickam

6431 Fannin Street,

MSB 7.102

Houston, TX 77030

**Uptake of LNPs into bEnd.3 brain endothelial cells and NSC-34 neuroblastoma cells**

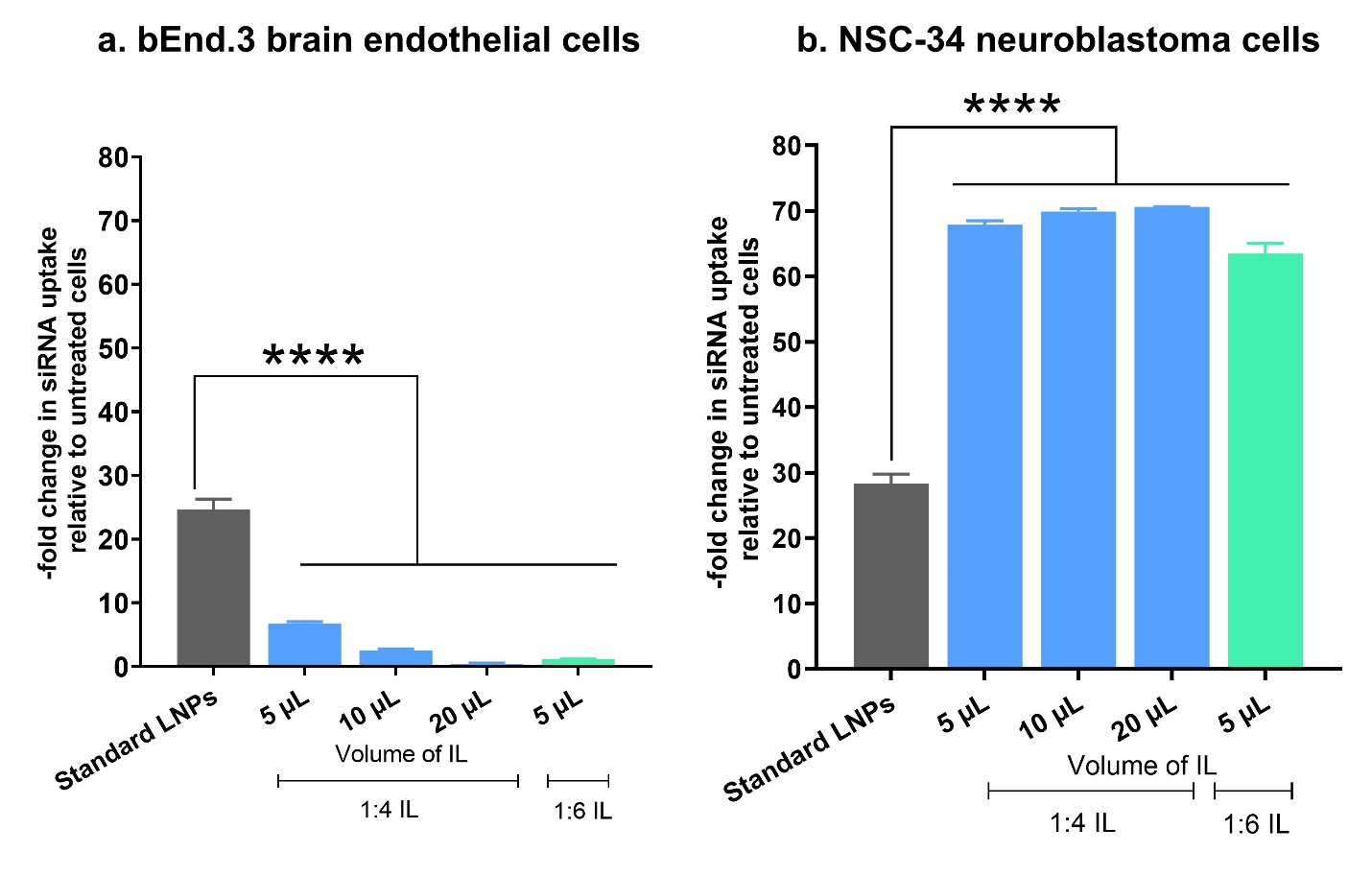

**Supplementary Figure 1. Cellular uptake of standard, 1:4 and 1:6 IL-coated LNPs containing Cy5 siRNA into bEnd.3 BECs and NSC-34 neuroblastoma cells using flow cytometry.** bEnd.3 BECs and NSC-34 neuroblastoma cells were treated for 4 h and 24 h respectively with standard, 1:4 and 1:6 IL-coated LNPs loaded with 50 nM Cy5 siRNA in complete growth medium in a humidified incubator at 37 °C and 5% CO_2_. Untreated cells were used as negative controls. Data are presented as a -fold increase in Cy5 siRNA uptake (% Cy5+ cells) relative to untreated cells (n=3). Statistical analysis was done using one-way ANOVA with Dunnett’s multiple comparisons test. ****p<0.0001.

**Gating strategy for RBC-hitchhiking experiment**

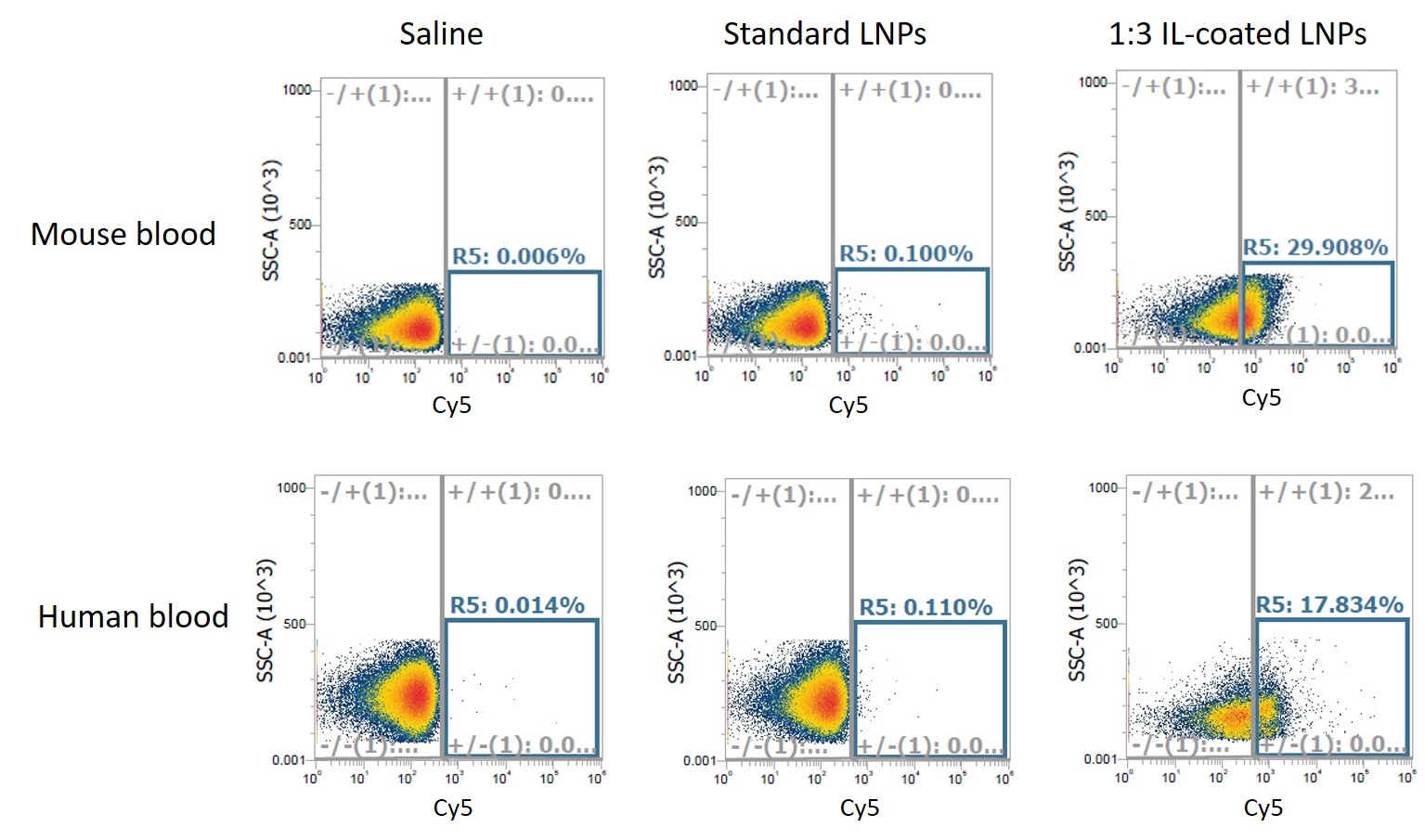

**Supplementary Figure 2. Gating strategy for flow cytometry.** Total LNPs hitchhiking on RBCs by flow cytometry for standard (uncoated) and 1:3 IL-coated LNPs. Cy5-siRNA-loaded LNPs were mixed with whole blood (1:3 v/v RBC: LNP), rotary-incubated at 37°C for 20 minutes followed by flow cytometry analysis to assess the fraction of RBC singlets with LNPs.
